# Nanovibrational stimulation of mesenchymal stem cells induces therapeutic reactive oxygen species and inflammation for 3D bone tissue engineering

**DOI:** 10.1101/2020.04.10.035568

**Authors:** Wich Orapiriyakul, Penelope M. Tsimbouri, Peter G. Childs, Paul Campsie, Julia Wells, Marc Fernandez-Yague, Karl Burgess, K. Elizabeth Tanner, Manlio Tassieri, R.M. Dominic Meek, Massimo Vassalli, Manus JP Biggs, Manuel Salmerón-Sánchez, Richard O.C. Oreffo, Stuart Reid, Matthew J Dalby

## Abstract

There is a pressing clinical need to develop cell-based bone therapies due to a lack of viable, autologous bone grafts and a growing demand for bone grafts in musculoskeletal surgery. Such therapies can be tissue engineered and cellular, such as osteoblasts combined with a material scaffold. Because mesenchymal stem cells (MSCs) are both available and fast growing compared to mature osteoblasts, therapies that utilise these progenitor cells are particularly promising. We have developed a nanovibrational bioreactor that can convert MSCs into bone-forming osteoblasts in 2D and 3D but the mechanisms involved in this osteoinduction process remain unclear. Here, to elucidate this mechanism, we use increasing vibrational amplitude, from 30 nm (N30) to 90 nm (N90) amplitudes at 1000 Hz, and assess MSC metabolite, gene and protein changes. These approaches reveal that dose-dependent changes occur in MSCs’ responses to increased vibrational amplitude, particularly in adhesion and mechanosensitive ion channel expression, and that energetic metabolic pathways are activated, leading to low-level reactive oxygen species (ROS) production and to low-level inflammation, as well as to ROS- and inflammation-balancing pathways. These events are analogous to those that occur in the natural bone-healing processes. We have also developed a tissue engineered MSC-laden scaffold designed using cells’ mechanical memory, driven by the stronger N90 stimulation. These new mechanistic insights and cell-scaffold design are underpinned by a process that is free of inductive chemicals.

---

Bone is the second-most grafted tissue after blood in humans and is used in a wide range of musculoskeletal surgeries.^1–3^ However, the use of autografts from patient donor sites is limited and has a high incidence of morbidity.^1–3^ Allograft is therefore widely used but also suffers from a lack of donor material, and is acellular and so lacks biological activity.^1–3^ Cellular therapies, used in combination with scaffolding materials, represent a future source of bone graft and regeneration approaches. As such, a growing number of mesenchymal stem cell (MSC)-based therapies are in clinical trial.^4^ We note that the term MSC is widely used and often refers to adherent stromal cells, as is typically the case in cell therapies.^5^ However, here we use the term MSC to more accurately refer to skeletal stem cells. These are a clonogenic population of non-hematopoietic bone marrow stromal cells that can recreate cartilage, bone, adipocytes and haematopoiesis-supporting stroma.^6, 7^

We have previously reported the development of a nanovibrational bioreactor that can stimulate MSC differentiation towards osteogenesis in 2D^8^ or 3D^9^ without the use of defined media, chemicals or highly specialised equipment (see supplementary Figure 1 for an image of the bioreactor). This approach offers significant advantages because it enables media, standard consumables, and MSC banks that have already been approved for use to be utilised. For example, in this study we used 6- and 24-well plates with our nanovibrational bioreactor by simply attaching them to a vibrating top-plate with magnets. The use of standard consumables also enables cells to be cultured in wells with scaffold materials, such as hydrogels, enabling us to think beyond just cell manufacture and towards tissue engineering.

The bioreactor employs the reverse piezo effect, in which a voltage is used to cause a mechanical expansion of a material, in this case a piezo active ceramic. The piezo ceramics are sandwiched between a large mass (aluminium block) and the ferrous top plate (which provides a magnetic interface).^10^ We have previously trialled a range of frequencies using this set up, with a fixed amplitude of 30 nm, and have found 1000 Hz to be optimal for MSC osteoinduction.^8^

Despite our ability to induce osteogenesis using this approach, we know little about how mechanistically nanovibrational mechanotransduction in MSCs induces osteogenic differentiation. We have previously reported altered cell adhesion and RhoA kinase (ROCK) activity in MSCs cultured in 2D in the nanovibrational bioreactor,^8^ and we have also previously identified TRPV1, transient receptor potential cation channel subfamily V member 1, as being implicated in 3D nanovibrational osteogenesis.^9^

We hypothesise that increasing nanovibrational amplitude will enhance osteogenic effect and related cell signalling. From our findings, we propose that increasing the nanovibrational amplitude that MSCs are exposed to exaggerates the underlying osteogenic mechanism, allowing us to further dissect MSC mechanotransduction mechanisms induced by nanovibration.

## Optimisation of 3D nanovibrational cultures

Type I collagen is a widely used 3D hydrogel scaffold, which we have previously used in nanovibrational experiments at 0.8 mg/ml concentration.^9^ An advantage of this collagen formulation is its low stiffness (E = *∼*26 Pa at 0.8 mg/ml, supplementary Figure 2A), which is well below the 30-40 kPa stiffness required to drive MSC osteogenesis^11, 12^ and its good biocompatibility as shown using alamar blue (supplementary Figure 2C). Importantly for nanovibrational stimulation, it adheres to the sides of cell culture plates, thereby providing mechanical integration with the plate. As a hydrogel, it is incompressible^13^. It thus acts as a solid volume when vibrated in a contained environment, such as the wells of a culture plate, providing good vibration propagation throughout its volume^9^ (Figure 1B). However, the action of cells within the collagen gel can induce the gel to contract from the edge of the well over long-term osteogenic cultures, which typically take 28 days to reach mineralisation.^14, 15^

**Figure 1.**
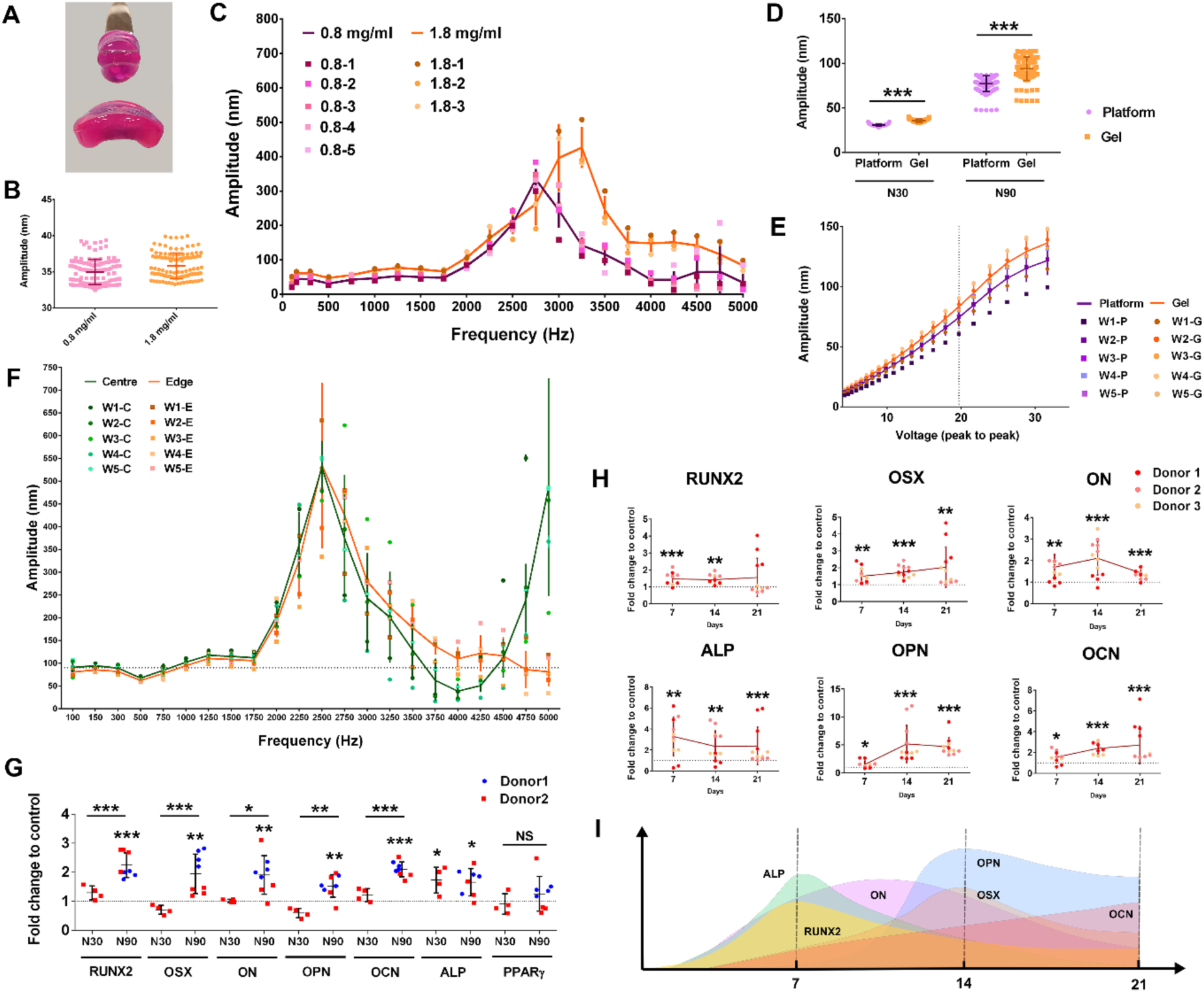
*3D MSC osteogenesis with N30 and N90 nanostimulation.* (A) Collagen gel constructed with 0.8 mg/ml (top) and 1.8 mg/ml (bottom) (note that both are front views and the gel diameter was 13 mm before removing from the well). (B) Interferometry showing that 0.8 mg/ml and 1.8 mg/ml collagen gels vibrate as expected with N30 stimulation (n=24). (C) Using interferometry, resonance effects were seen at frequencies >2000 Hz, but no resonance peaks were seen at 1000 Hz, for both 0.8 mg/ml and 1.8 mg/ml collagen gels (n=3-5). (D) Interferometry results for N30 and N90 nanostimulation, showing good fidelity of vibration in the 1.8 mg/ml collagen gel (n=24). (E) Interferometry showed a linear voltage – amplitude relationship for both the vibration platform and the 1.8 mg/ml collagen gel (n=5) between 12 and 27 Vpp. (F) No resonance frequencies were seen at 1000 Hz at either the centre or edge of the 1.8 mg/ml collagen gels (n=5). (G) Osteogenic marker gene expression in MSCs, as assessed by qPCR for N90 compared to N30 (or control), after 9 days of culture in 3D nanovibrational stimulation conditions in 1.8 mg/ml collagen gels. Osteogenic marker gene expression was enhanced in N90 conditions, compared to N30 or control conditions (d=2, r=4, t= 3). (H) Osteogenic transcript expression in N90 conditions in 1.8 mg/ml collagen gels at days 7, 14 and 21 of culture. (I) Schematic of expression maxima with time (d=3, r=4, t= 3). Significance calculated using ANOVA with Tukey multiple comparison, where *=p<0.05, **=p<0.01 and ***=p<0.001. Error bars represent means±SD. The data shows good fidelity and improved efficacy of 3D nanostimulation with change in gel stiffness and with increased vibration amplitude. Abbreviations, d = number of donors assessed; r = number of wells tested; t = technical replicates. Donors are MSCs derived from different donor sources.

In order to overcome this issue, we trialled a 1.8 mg/ml collagen gel. While this gel formulation is stiffer (E = *∼*161 Pa, supplementary Figure 2A), it remains significantly below the stiffness required to induce MSC osteogenesis (>20 kPa).^11, 12^ It was also easier to handle (as shown in Figure 1A), which could have positive implications in the operating theatre, where surgeons need to remove a cell product from a dish to place into a patient. Importantly, vibrational fidelity was similar to that of the 0.8 mg/ml gel, with a 30 nm displacement at the vibration plate surface inducing 35 nm vibrations in both the 0.8 and 1.8 mg/ml gels, as indicated by laser interferometry (Figure 1B). Furthermore, little evidence of resonance effect was seen in any replicates of the 0.8 and 1.8 mg/ml gels at 1000 Hz driving frequency (Figure 1C). By looking at collagen-plastic detachment over a longer culture period, we found that while the 0.8 mg/ml gels contracted within a 30 day culture period, with typical cell seeding of 40,000 MSCs/ml collagen (supplementary Figure 2B), the 1.8 mg/ml gels did not contract until >70 days when seeded with either 40,000 or 80,000 cells/ml and exposed to 1000 Hz, 30 nm vibrational stimulation. This time period extends way beyond the usual duration of our MSC nanovibration experiments (supplementary Figure 2B).

A key aim of this research was to investigate if larger amplitude can induce more pronounced changes to allow us to infer cell mechanism with more clarity. To investigate this, we selected a second amplitude of 90 nm, which we used in addition to a previously tested amplitude of 30 nm (referred to as N30 and N90, respectively). For both gels, we observed a slight increase in amplitude, relative to that of the top plate, from 30 to *∼*35 nm with N30, and from 90 to *∼*100 nm with N90 (Figure 1D). We note that the voltage-amplitude relationship was linear between 12 and 27 Vpp (voltage peak-to-peak, the region in which we operate) in the 1.8 mg/ml collagen gel, as it was when measured on the bioreactor top plate (Figure 1E), again demonstrating the fidelity of the system.

We next assessed resonance using interferometry. At 1000 Hz, the bioreactor generated reliable displacements without resonance problems at the edge and the centre of the 1.8 mg/ml hydrogels (Figure 1F). Osteogenic marker expression was also assessed by qPCR. After 9 days of MSC stimulation in 3D, using 1.8 mg/ml gels and N30 stimulation, alkaline phosphatase (ALP) expression was detected (Figure 1G). However, osteogenic stimulation was significantly more pronounced following N90 stimulation, with several osteoblast markers expressed after 9 days, including ALP, runt related transcription factor 2 (RUNX2), osterix (OSX), osteonectin (ON), osteopontin (OPN) and osteocalcin (OCN) (Figure 1G). PPAR*γ* (peroxisome proliferator activated receptor *γ*) expression, an adipocyte marker, was also assessed to gauge whether the nanovibrational effect was osteospecific. No induction of PPAR*γ* was observed with N30 or N90 stimulation at 9 days of culture (Figure 1G). Alamar blue staining showed that there were no cytotoxic effects of N30 or N90 stimulation in the 1.8 mg/ml collagen gels or in the control conditions with/without osteogenic media (supplementary Figure 2C).

## Higher amplitude stimulation increases osteogenesis and ion channel expression

We next assessed how nanostimulation at N30 and N90 conditions affected the expression of adhesions and ion channels that have been previously implicated in N30-stimulated MSC osteogenesis.^9^ To do so, we used a protein array containing a range of receptors and channels. Using this array, we observed the upregulated expression of beta 1, 3 and 5 integrins after 9 days of MSC culture in both N30 and N90 conditions (Figure 2A). These integrins function as receptors for a wide range of extracellular matrix (ECM) proteins. A range of collagens were also upregulated in N30- and N90-stimulated MSCs (Figure 2A); we note that integrin beta 1 is used by cells to attach to collagens.^16^ Bone morphogenetic protein 2 (BMP2) receptor BMPR1 was also upregulated following nanovibrational stimulation (Figure 2A). Most interestingly, however, was the pattern of ion channel expression by MSCs. In agreement with previous results,^9^ N30 conditions stimulated the expression of TRPV1 (transient receptor potential cation channel subfamily V member 1) (Figure 2A). However, under N90 conditions, TRPV1 was more highly expressed, as were TRPA1, Piezo1 and 2, and KCNK2 (potassium channel subfamily K member 2) (Figure 2A), each of which are mechanosensitive ion channels that can be opened by stretch, for example by membrane deformation^17, 18^, or by myosin contracting the cytoskeleton^19, 20^. They also reportedly transduce high frequency vibrational forces, such as in the ear^21, 22^ (we note that 1000 Hz is in the audible range). Moreover, KCNK2 (also known as TREK1) has been associated with low-frequency osteoinduction of MSCs via magnetic twisting cytometry.^23^

**Figure 2.**
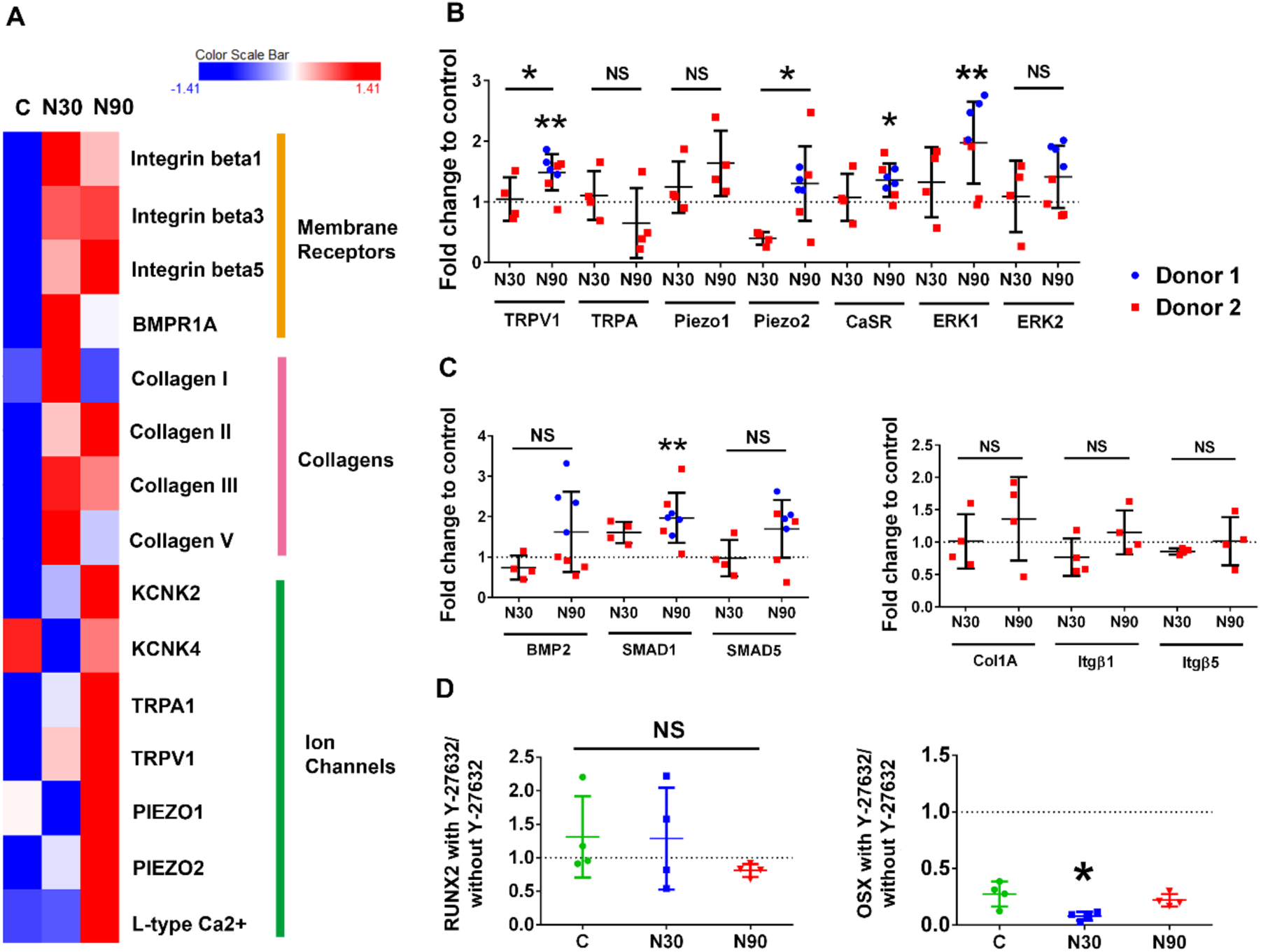
*Adhesion and ion channel expression by MSCs when cultured with vibrational amplitude of N30 or N90.* (A) Protein array data, presented as a heatmap. It shows the expression levels of a range of adhesion, extracellular, and ion channel proteins in MSCs cultured in control conditions (C) and in conditions of N30 and N90 vibrational amplitude, after 9 days of stimulation (d=2, r=4, t= 3). Red denotes increased expression, blue denotes reduced expression, relative to control. (B, C) Ion channel expression (B), BMP family member’s expression (C, left) and adhesion molecule expression (C, right), as assessed by qPCR, for control, N30 and N90 cultures after 9 days of stimulation (d=2, r=4, t= 3). (D) qPCR analysis of RUNX2 and OSX expression, with or without ROCK inhibition (Y-27632), for control, N30 and N90 cultures after 9 days of stimulation (d=1, r=4, t= 3). Significance calculated using ANOVA with Tukey multiple comparison where *=p<0.05 and **=p<0.01. Error bars represent means±SD. Data shows that ion channel, and extracellular matrix and adhesion proteins are more highly expressed in N90 conditions relative to N30 conditions and that inhibiting ROCK has only a small effect on osteogenesis. Abbreviations, d = number of donors assessed; r = number of wells tested; t = technical replicates.

It is notable that Piezo1 is down-regulated in N30 conditions and up-regulated in N90 conditions, compared to control. This might be a time-dependent phenomenon or a gated phenomenon, wherein higher levels of stimuli are more likely to activate threshold-dependent mechanisms. Indeed, the force-dependent activation of piezo1 has been compared to a switch;^24^ for example, Piezo1 has been linked to ATP signalling in MSCs in a threshold-dependent manner.^25^

We assessed protein levels with the protein array and transcript levels with qPCR at the same time points. Ion channels showed less change at the transcript level than at the protein level, with N90 stimulation producing the greatest upregulation of TRPV1, Piezo2 and CaSR (calcium sensing receptor) expression, and of the downstream target, ERK1 (extracellular signal related kinase 1). Among the assessed BMP signalling family members, only the expression of SMAD1 (small mothers against decapentaplegic 1) was induced by N90 stimulation (Figure 2C, left), while the assessed collagen genes were not expressed by MSCs cultured with N30 or N90 stimulation (Figure 2C, right). Inhibiting cytoskeletal tension with the ROCK inhibitor, Y-27632, produced only a subtle loss of osteogenesis, with only N30 conditions indicating that intracellular tension is important for cell responses to 3D nanovibrational stimulation (Figure 2D).

From these results, we propose that vibrational amplitude at N90 provides a more powerful osteogenic cue than does N30, and that ion channel expression is particularly increased by this higher amplitude.

## Reactive oxygen species and nanovibrational stimulation

To investigate further the cellular pathways that are altered by nanovibrational stimulation in MSCs, we took an untargeted metabolomics approach. Cells were lysed after culture for 1 or 2 weeks in control, N30 or N90 conditions, and then analysed by liquid chromatography (LC) – orbitrap mass spectrometry (MS).^26^ Heatmap analysis and principle component analysis (PCA) revealed that lipids are the largest differentially regulated metabolite group (Figure 3A) and that the metabolome of N30 and N90 cultured cells MSCs diverged from each other and the control group (Figure 3B).

**Figure 3.**
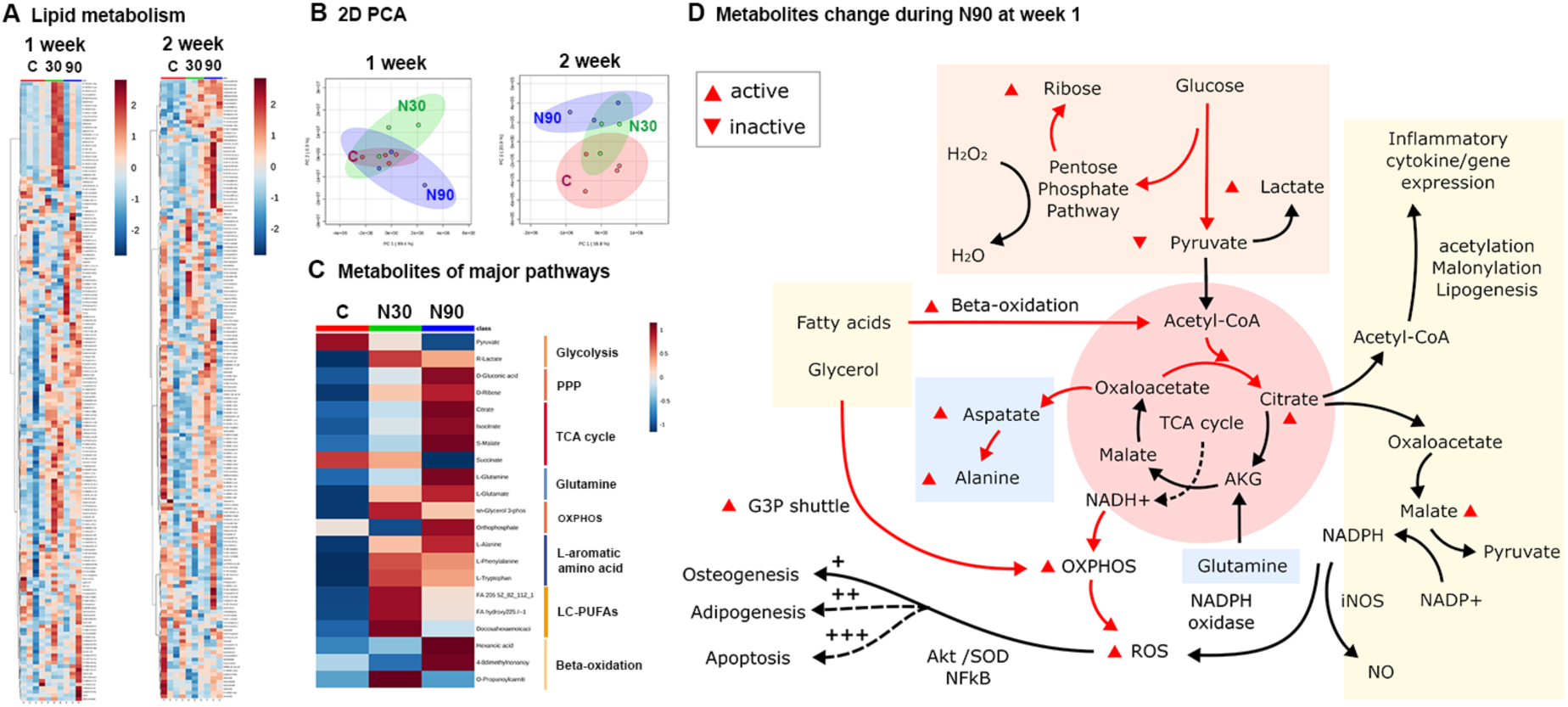
*Untargeted metabolomic analysis of MSCs cultured in N30 and N90 stimulation conditions.* (A) Lipid heatmaps of MSCs after 1 and 2 weeks of nanostimulation at N30 and N90 amplitudes. (B) Principle component analysis (PCA) of MSC lipid data after 1 and 2 weeks of culture in N30 and N90 nanostimulation conditions, compared to control. (C) Observed metabolite changes in ROS pathways following 1 week of culture under N30 or N90 conditions. (D) Schematic of potential pathways derived from the heatmap data (d=1, r=4, t=1). The data indicate the activation of ROS and redox-balancing pathway occurs in MSCs cultured in conditions of increasing nanostimulation amplitude. Abbreviations, d = number of donors assessed; r = number of wells tested; t = technical replicates.

After 1 week of culture, the metabolites of major respiration-related pathways, including glycolysis, the pentose phosphate pathway, TCA cycle, oxidative phosphorylation (OXPHOS), L-aromatic amino acid, long-chain polyunsaturated fatty acids (LC-PUFAs) and *β*-oxidation) were typically up-regulated to differing levels by N30 and N90 conditions (Figure 3C). N90 conditions produced the most differentially regulated responses relative to unstimulated control conditions. The responses to N90 were therefore used to build a pathway map. The pathways that were affected include inflammation and reactive oxygen species (ROS) and the pentose phosphate pathway (PPP), which acts to balance oxidative stress^27^ (Figure 3D). Together, these data indicate that nanovibrational stimulation triggers an energetic response in cells, as evidenced by increased glycolytic and tri-carboxylic acid (TCA) cycle metabolite levels^28^ (Figure 3D). Metabolite pathways were also analysed using Ingenuity pathway analysis (IPA). The IPA analysis further supported these results. After 1 week of nanovibrational stimulation, metabolic pathways were up-regulated in both N30 and N90 conditions, relative to control conditions, with greater up-regulation seen in N90 conditions (supplementary Figure 3). By 2 weeks of nanostimulation, pathways were mostly down-regulated in N30 conditions, but remained predominantly up-regulated in N90 conditions (supplementary Figure 4). This, again, suggests that MSC osteogenesis stimulated by nanovibration is an energetic process and that the greater the stimulus, the greater the observed effects on the cell. We also note, from increases in ROS- and PPP-related metabolites, that redox balancing might also be potentially occurring to counter oxidative stress (Figure 3D).

To follow up the observation that redox balancing potentially occurs in response to nanostimulation, we again used IPA to analyse the metabolomic data. By looking at oxidative stress (Figure 4A), we observe pathways being increasingly activated in N90 relative to N30 conditions. Looking at the individual metabolites inferred as contributing towards oxidative stress, we observe the same metabolites forming the network, but being more highly activated in N90 conditions (Figure 4B, C). We next used 2’-7 dichlorodihydrofluorescein diacetate (DCF-DA) flow cytometry to measure ROS. While a small increase was noted in N30 conditions, relative to the control, this increase became statistically significant in N90 conditions (Figure 4D).

**Figure 4.**
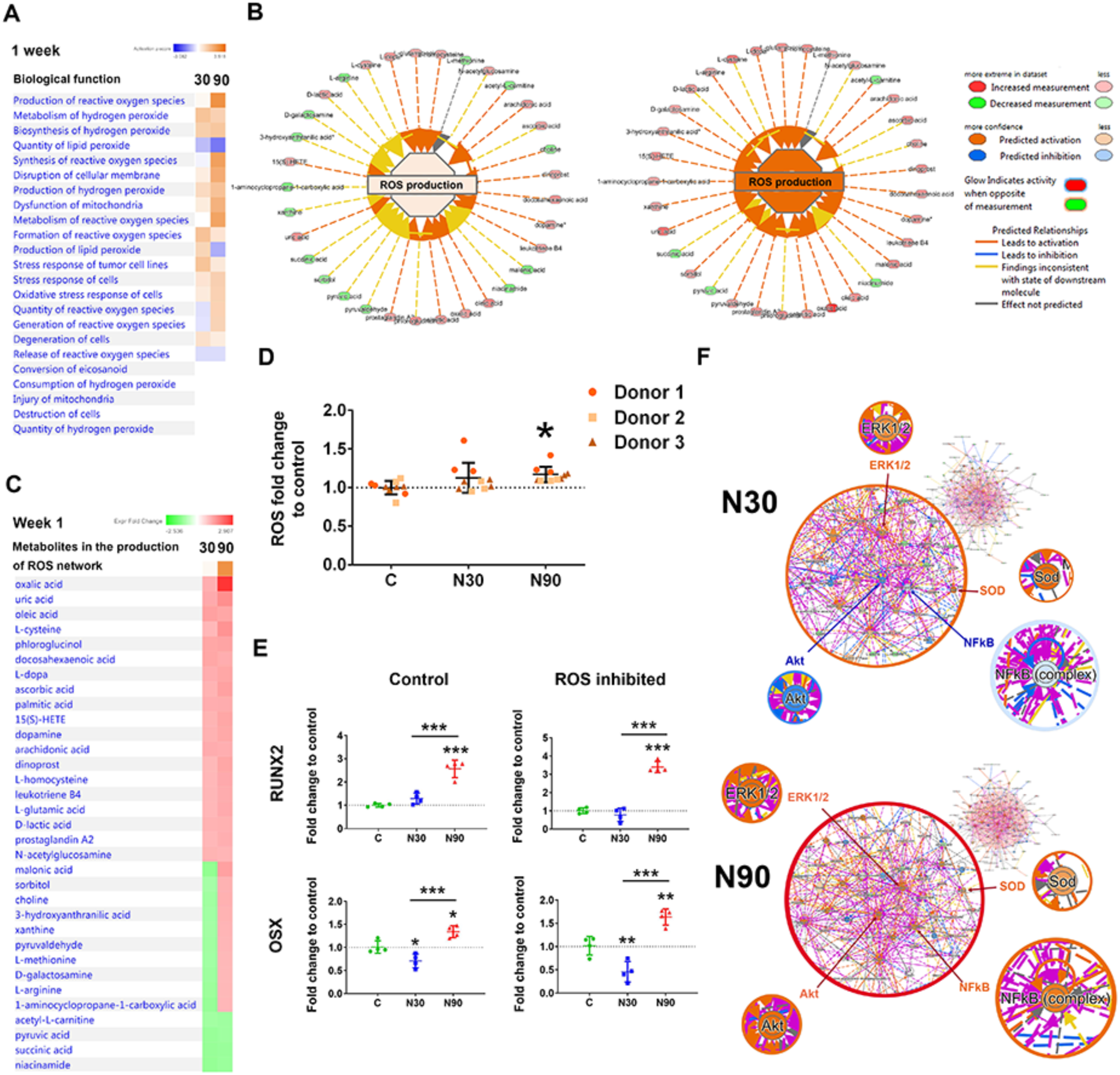
*Metabolomic analysis of MSCs cultured with N30 and N90 nanostimulation.* (A) Observation of ROS based pathways. (B, C) Metabolites involved in ROS after MSCs were cultured for 7 days of under control, N30 and N90 conditions (d=1, r=4, t=1). (D) Dichlorodihydrofluorescein diacetate (DCF-DA) flow analysis, showing that ROS levels increase with nanostimulation amplitude, reaching significance following MSC stimulation at N90 (d=3, r=3, t=1). (E) qPCR analysis of *RUNX2* and *OSX* expression in MSCs after 9 days of culture under N30 or N90 conditions. Both markers were upregulated following N90 stimulation in this donor cell line, with ROS inhibition having little effect on their expression (d=1, r=4, t=3). (F) (top) Ingenuity pathway analysis of metabolite networks induced by N30 stimulation, showing the predicted up-regulation of ERK1/2 (osteogenic commitment) and SOD (ROS) and the predicted repression of Akt and NF*κ*B (REDOX balancing). (bottom) N90 stimulation is predicted to result in the activation of all these pathways (d=1, r=4, t=1). For D and E, error bars represent means±SD. Significance calculated using ANOVA with Tukey multiple comparisons, where *=p<0.05, **=p<0.01 and ***=p<0.001. The data indicate that MSCs generate ROS while committing to osteogenesis and that, when the signal becomes stronger, the cells also activate REDOX balancing pathways. They also indicate that ROS itself is not a driver of osteogenesis. Abbreviations, d = number of donors assessed; r = number of wells tested; t = technical replicates.

Previous studies have linked small increases in ROS to enhanced osteogenesis; however, large increases in ROS are also linked to the suppression of osteogenesis.^29–31^ To investigate this issue in our experimental system, we used N-acetyl cysteine to inhibit ROS (Figure 4E) and then assessed *RUNX2* and OSX expression in MSCs after 9 days of culture by qPCR. Our results show that while RUNX2 and OSX were expressed at a low level in N30 conditions, both markers were significantly up-regulated in N90 conditions, again showing that enhanced osteogenesis occurs with the larger amplitude. In addition, little change in marker expression was seen in response to ROS inhibition, indicating that ROS do not have a detrimental effect on osteogenesis and are a likely by product, rather than a driver, of osteogenesis.

Using IPA activity predictor, where metabolite networks are linked to biochemical signalling hubs, some consistencies and some similarities and some differences in signalling could be observed between N30 and N90 stimulation (Figure 4F). ERK 1/2 is predicted to be up-regulated, as are the superoxide dismutase (SOD) pathways for both N30 and N90 stimulation. ERK 1/2 stimulation is widely reported to be important for MSC osteogenesis, as it is implicated in the phosphorylation and activation of RUNX2, the osteogenic master transcription factor.^32–34^ SOD is used by cells to counter balance the effects of ROS^35^, and so this finding fits with our observations of increased ROS production and PPP activation (Figure 4 B, D). Interestingly, the results of stimulation at N90 also implicate the NF*κ*B (nuclear factor kappa-light-chain-enhancer of activated B cells) and Akt (protein kinase B) pathways, which are involved in cell survival and in preventing apoptosis, and are both linked to antioxidant function.^36, 37^

## Inflammation and nanovibrational stimulation

Increased levels of ROS lead to inflammation^38^, and a small degree of inflammation is implicated in the natural bone healing process, while high levels of inflammation prevent osteogenesis.^39^ Given this, we hypothesised that inflammation might be observed in MSCs following their culture under N90 conditions. To explore this, we assayed the expression of the inflammatory markers, interleukin 6 (IL-6), NF*κ*B, and tumour necrosis factor (TNF*α*), by qPCR after 9 days of N90 culture, as well as that of two mitogen activated protein kinases (MAPKs), ERK (which is involved in cell proliferation and osteogenesis^32–34^) and of c-jun n-terminal kinase (JNK, which is also implicated in osteogenesis^32^ but is better known for being activated by ROS or inflammation to mediate cytokines and apoptosis).^40^ We observed increased expression of TNF*α*, ERK and JNK after 9 days of culture and N90 stimulation compared to unstimulated control (Figure 5A).

**Figure 5.**
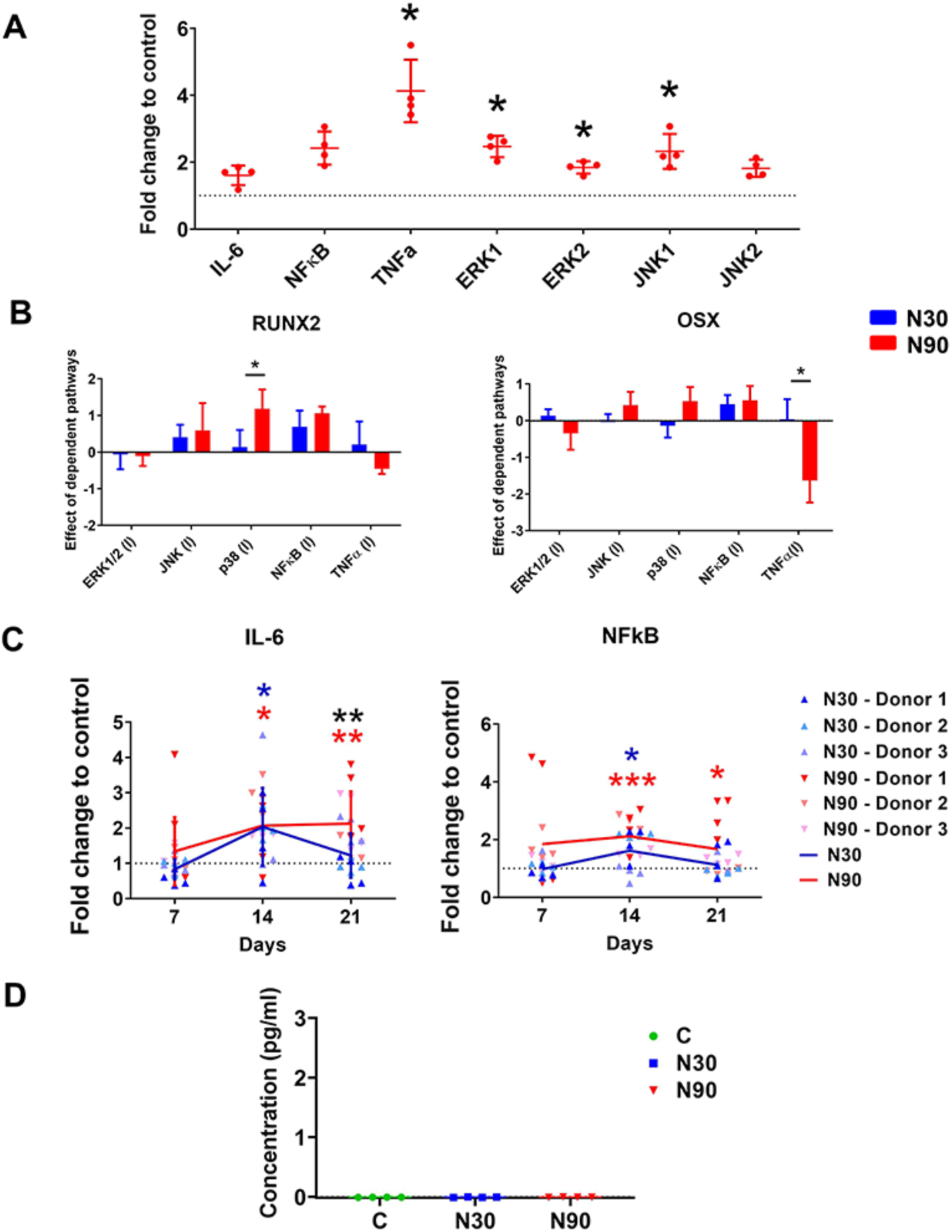
*Nanostimulation at N90 upregulates inflammatory markers in MSCs.* (A) Increased expression of *TNFα*, *ERK1/2* and *JNK1* in MSCs cultured with N90 stimulation for 9 days, as assessed by qPCR (d=1, r=4, t=3). (B) Inhibition studies with qPRC for RUNX2 and osterix (OSX) showed that p38 MAPK inhibited RUNX2 expression and TNF*α* enhanced OSX expression with N90 stimulation compared to N30 (d=1, r=4, t=3). (C) IL-6 and NF*κ*B show increased expression to day 14 of culture, and decreased expression to day 21, as assessed by qPCR; increased expression was more persistent for N90 stimulation (d=3, r=4, t=3). (D) IL-1 showed no detectable expression with either N30 or N90 stimulation, as assessed by ELISA (d=1, r=4, t=1). Error bars represent means±SD, significance calculated using ANOVA with Tukey multiple comparison, where *=p<0.05, **=p<0.01 and ***=p<0.001 (note that blue and red asterix show the significant difference of N30 and N90 to control, while black asterix represent significant difference between N30 and N90). The data shows a very low-level inflammatory response that is not detectable at the protein level. Abbreviations, d = number of donors assessed; r = number of wells tested; t = technical replicates.

We next inhibited these inflammatory pathways and p38 MAPK (which is activated by cell stress and is involved in apoptosis and differentiation control^41^), and assessed the expression of the osteogenic markers, RUNX2 and OSX, in MSCs cultured under N30 and N90 stimulation conditions. It was seen that compared to N30, N90 osteogenesis (RUNX2) was enhanced by p38 MAPK inhibition and reduced (OSX) by TNF*α* inhibition (Figure 5B).

Looking at the pro-inflammatory cytokine (IL-6)^42^ and the inflammation response factor (NF*κ*B)^36^ at days 7, 14 and 21 of MSC culture, we observed that the expression of these inflammatory mediators tracked each other; their expression increased to day 14 and then reduced (Figure 5C), most notably with N90. This suggests that nanostimulation induces an inflammatory response that is then countered by the cells, and that the magnitude of this response scales with amplitude. We used ELISA to assess the levels of the major proinflammatory cytokine, IL-1*β*^43^ and found that it was undetectable at N30 or N90 conditions, with concentrations below the sensitivity of the standard curve (Figure 5D). This demonstrates that the inflammatory response we observed is small rather than constituting real inflammation, at a level more likely to be positive in terms of bone formation.^39^

Our unpublished data suggest that MSCs from ∼1 in 20 donor samples do not respond to nanovibrational stimulation. We looked at NF*κ*B, which is linked to osteogenesis in MSCs, from one of these donors. In this donor, there was little evidence of osteogenesis having occurred after N30 or N90 stimulation, based on the expression of RUNX2, OSX and OPN after 9 days of culture, (supplementary Figure 5A). However, when NF*κ*B was inhibited with TPCA-1, the expected increase in these osteogenic markers was observed (supplementary Figure 5A). Furthermore, nanovibration resulted in the decreased expression of NF*κ*B in direct proportion to vibration amplitude (supplementary Figure 5B). We thus speculate that the inflammatory background of the donor and/or dysregulation of NF*κ*B might increase or decrease the osteogenic capacity of individual MSCs.

## Tissue engineering using nanovibration

These new data demonstrate that increasing nanoscale amplitude can enhance osteogenesis through a low-level ROS/inflammatory axis in 3D MSC cultures. Here, we look to see if these changes hold as we develop a composite more suitable to being used for tissue engineering.

The gels used in this report are very soft (*∼*26-161 Pa) and thus problematic to handle. Exploiting this osteogenic effect in a more structurally stable scaffold is of vital importance for the clinical translation of this technology. However, there are some constraints. The scaffold needs to be physically integrated with the well plate, and thus collagen gels are useful as they are biocompatible and attach to the sides and bottom of culture dishes (unless the cells detach the gel through contraction). Collagen gels are also highly hydrated. Water is incompressible, meaning that when constrained, such as in a culture plate well, it acts as a solid object;^13^ hydrogels are mainly water and this means that the cells experience vibration in all parts of the gel as we demonstrate on top of the gels using interferometry (Figure 1D).

We thus decided to generate a composite gel using a collagen sponge to provide rigidity while maintaining biocompatibility. Insoluble collagen was freeze dried to form a highly porous structure (pore size 227.74±72.93, measured using Feret’s diameter^44^), which was *∼*4 mm high and *∼*11 mm in diameter (Figure 6A), and has an elastic modulus of 1.08±0.29 MPa (Zwick-Roell compression testing). The acellular scaffolds were held in place with a small weight while 2.5 mls of MSC-containing neutralised collagen solution was poured over them. This sets the scaffolds within the wells of a 12-well plate and provides a 3D MSC source (Figure 6B). Once the gel is set and the scaffold set in place, the weight is removed and a small amount of fresh collagen is added to complete the gel (Figure 6B).

**Figure 6.**
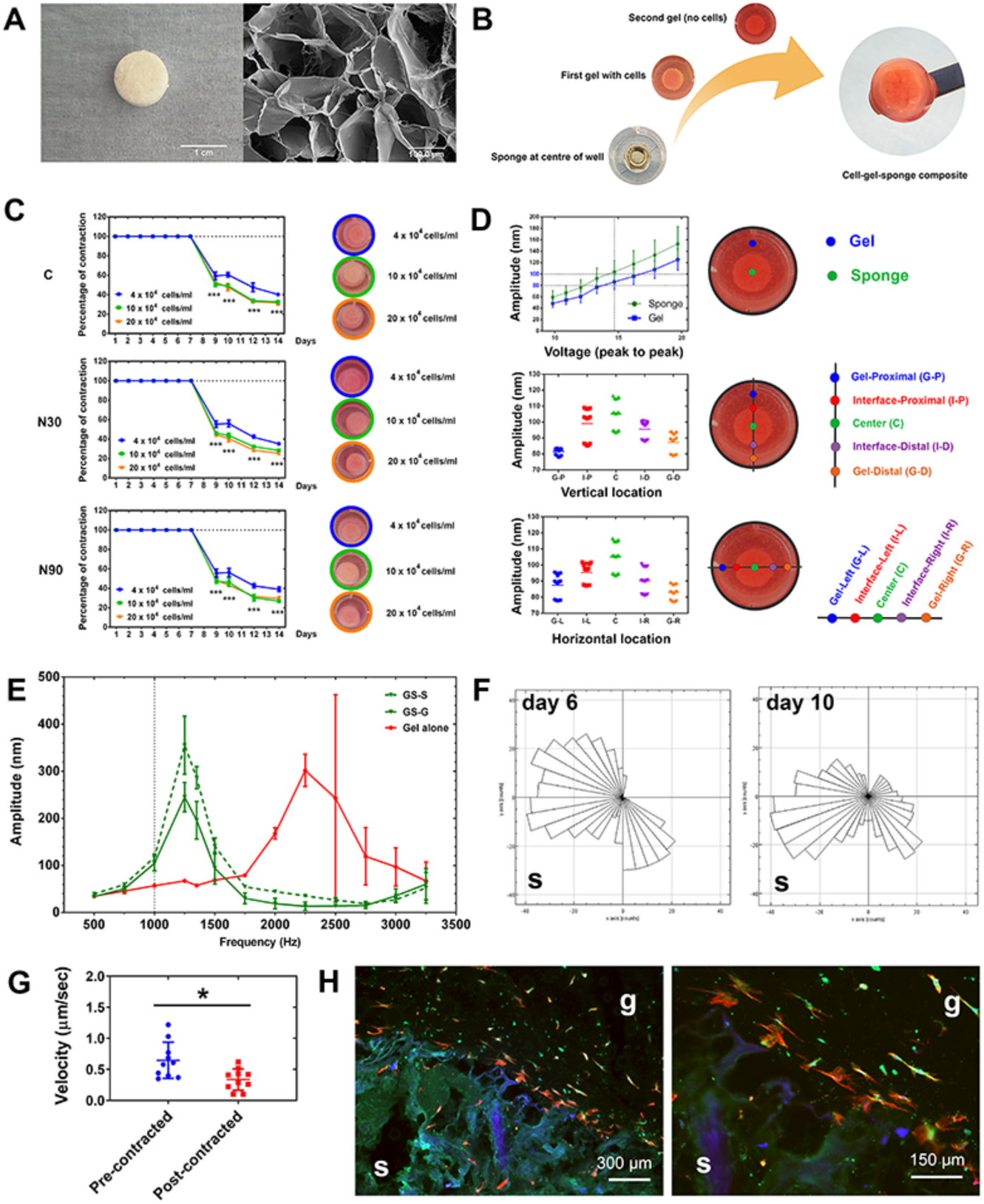
*Design of a gel-sponge composite for nanovibrational stimulation.* (A) A freeze-dried collagen sponge (left) showing its pore structure by SEM (right). (B) Schematic of the composite gel’s fabrication. The sponge is held down with a weight while the cell/gel mixture is poured over it and sets. After the weight is removed, more collagen is poured on. At a user-defined time, the cell-containing collagen gel can be released from the well edges and allowed to contract onto the sponge to form an easy to handle construct. (C) Gel constructs, containing 4, 10 and 20×10^4^ MSCs/ml, were stimulated for 7 days, and then the gels detached from the well. All gels contracted onto the sponge within 2 days, and the sponge prevented further contraction (d=1, r=4, t=1). (D) Interferometry testing of vibrational transmission in the composite gel. A linear voltage amplitude relationship was seen both at the sponge and gel positions (top). Looking at points across the gel-sponge composites (middle and bottom), slightly higher amplitudes were observed at the sponge position compared to the gel at N90 stimulation, with up to 20 nm variance observed (n=3-6). (E) Increasing vibration frequencies were assessed using interferometry. The sponge induced resonant frequencies at ∼1250 Hz. At 1000 Hz, however, little evidence of resonant effects was observed (n=3, GS; gelsponge composite, G; gel area, S; sponge area). (F) MSC migration in the gel towards the sponge, shown at day 6 (pre-contraction) and at day 10 (post contraction). Movement towards the sponge increased with contraction (S= sponge location). (G) Cell velocity, however, decreased post-contraction (d=1, r=1, t=10). (H) Histology of gel-sponge composite sections showing MSC migration into the gels (red=10, green=10, blue=10; g=gel, s=sponge). The data show that this fabricated gel-sponge composite can facilitate cell migration and the application of nanovibrational stimulation. Abbreviations, d = number of donors assessed; r = number of wells tested; t = technical replicates.

In order to make scaffolds that are easier to handle for potential clinical applications, we utilised the adherent properties of collagen to allow for vibration fidelity. We did so by vibrating the gels for a period and then allowing the contractile properties of the cells in the collagen to pull the gels onto the sponge. To test this, gels were seeded with 4, 10 and 20×10^4^ cells per ml of MSCs and cultured with/without N30 and N90 stimulation for 7 days. Gels were then released from the sides of the wells and contraction observed; all gels in all conditions contracted on to the sponge within 2 days, with 10×10^4^ and 20×10^4^ MSCs contracting the gels more than 4×10^4^ MSCs (Figure 6C). Next, interferometry was used to observe gel vibrational response to N90 input. We observed that vibration was higher in the well centre (over the gel) at just over 100 nm and was lower at the well edges at 80-90 nm (Figure 6D); acceptable vibration fidelity was seen.

Using N90 conditions with 4 × 10^4^ MSCs/ml, we looked for resonant frequency at 1000 Hz. While measuring at the centre (sponge, S) and edge (gel, G) of the gel-sponge composite, no resonance effects were seen at increased amplitude of just over 100 nm (Figure 6E), enabling us to proceed with 1000 Hz stimulation.

We also looked at MSC migration into the gel composite at 4 × 10^4^ MSCs/ml without N90 stimulation. MSC migration, while always towards the sponge, became more targeted post-contraction (day 10) compared to pre-contraction (day 6) (Figure 6F). The velocity of MSC migration, however, decreased post contraction (Figure 6G). Histology at day 12 confirmed that cell migration into the gels had occurred (Figure 6H).

We next moved to consider osteogenesis and also mechanical memory (or mechanical priming) of MSCs in the gel-sponge composites. This was in order to check firstly that the composites could be used with nanovibrational stimulation and then secondly to see if the composites could be vibrated for shorter-term cultures with prolonged osteogenesis (i.e. with memory of the initial vibration). Thus, in the gel-sponge composites, 4 × 10^4^ MSCs/ml with/without N30 and N90 stimulation were assessed for osteogenic markers by qPCR after 1, 2 and 3 weeks of vibration without gel detachment. We observed the increased expression of pro-osteogenic markers at both N30 and N90 stimulation, and this increase was greater at N90 stimulation (Figure 7A). The expression of IL-6 and NF*κ*B were also tested to assess the inflammatory response of MSCs during nanostimulation in the composite. Although IL-6 and NF*κ*B were initially expressed, their expression quickly reduced to background levels (Figure 7B), concurring with our low-level inflammation-osteogenesis hypothesis (Figure 7C).

**Figure 7.**
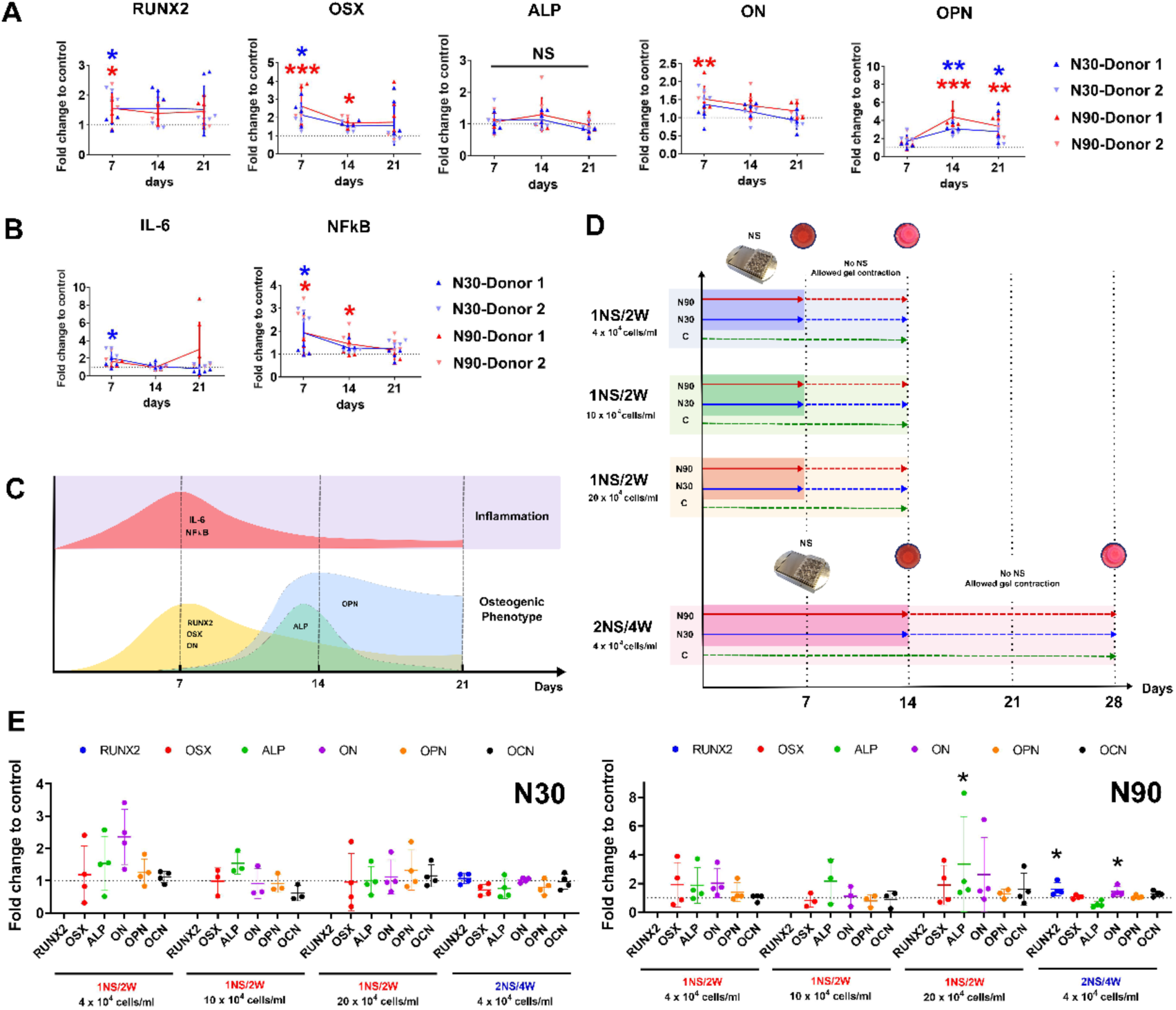
*Osteogenesis and mechanical memory of gel-sponge constructs.* (A) Osteogenic marker expression analysis by qPCR of MSCs cultured for 1, 2 and 3 weeks in the composite gel without gel detachment. (d=2, r=4, t=3). (B) IL-6 and NF*κ*B inflammatory marker expression by qPCR of MSCs cultured for 1, 2 and 3 weeks in the composite gel without gel detachment. The markers are initially up-regulated and then rapidly return to control levels (d=2, r=4, t=3). (C) Schematic of the expression profiles of inflammatory and osteogenic markers in MSCs cultured under N90 stimulation. (D) Schematic of the stimulation, detachment and experimental termination regimens for mechanical memory analysis. They were as follows: 1 week of nanostimulation followed by gel detachment from the sides of the well, 1 week of contraction (1NS/2W) for 4, 10 and 20×10^4^ MSCs per ml of collagen, using N30 and N90, and 2 weeks of nanostimulation followed by gel detachment from the sides of the well; and 2 weeks of contraction (2NS/4W) for 4×10^4^ MSCs per ml of collagen using N30 and N90. (E) Using N30, very little evidence of mechanical memory post cessation of nanostimulation and subsequent contraction was observed. However, following N90 stimulation, evidence of mechanical memory was seen using 20×10^4^ MSCs per ml for the 1NS/2W regime with enhanced ALP expression. Mechanical memory was also observed using N90 for the 2NS/4W regime with 4×10^4^ MSCs per ml with enhanced expression of *RUNX2* and *ON* (d=1, r=4, t=3). Error bars represent means±SD, significance calculated using ANOVA with Tukey multiple comparison where *=p<0.05, **=p<0.01 and ***=p<0.001 (note that where applicable, blue and red asterix show the significant difference of N30 and N90 to control while black asterix represents significant difference between N30 and N90). The data show that MSC osteo-differentiation occurs within the composite constructs and that, with N90, mechanical memory can be used to enable the production of a contracted, easy to handle, tissue-engineered product.

Finally, we assessed two different culture regimes for contracting the gel onto the sponge to enable the manufacture of free-floating scaffolds with good handling properties. The first used 1 week nanostimulation before gel detachment (osteogenic markers were upregulated in the MSCs in the composites) (Figure 7A) to produce free floating cell-composite scaffolds. Another week of culture was then allowed to promote full contraction onto the scaffold, giving a 2 week production timeline (1:1); this was performed with 4, 10 and 20×10^4^ MSCs per ml of collagen. The second regime used 2 weeks of nanostimulation before gel detachment and a further 2 weeks of culture (2:2), giving a 4-week production timeline for the free-floating cell composites. For N30, very little osteogenesis was observed with either the 1:1 or 2:2 regimes (figure 7E). However, with N90, osteogenic transcriptional changes were observed for the 1:1 regime that used the highest level of cell loading (20×10^4^), and with the longer 2:2 regime that used the lower cell loading level (4×10^4^) (Figure 7E).

## Summary

By increasing the amplitude of nanostimulation, we have been able to better dissect the cellular mechanisms of nanovibrational osteogenesis in MSCs. Many 2D osteogenesis studies have highlighted the central role of intracellular tension. In this 3D study, we can see clear changes in the regulation of integrin receptors and of ECM components, such as collagen (Figure 2A). We also observed the differential expression of known integrin-linked osteogenic pathways, such as ERK (Figure 2B). In all cases, stimulating MSCs with N90 produced a more marked transcriptional and protein-level response. However, reducing intracellular tension via ROCK inhibition did not significantly impact osteogenesis in our system (Figure 2D). Our results thus suggest that while adhesion is modulated and contributes to osteogenesis, it is not the central driver.

Several ion channels were also up-regulated during nanovibrationally stimulated MSC osteogenesis. Ion channel expression, and the number of channels showing enhanced expression, were linked to increasing amplitude (Figure 2A, B). For example, at the protein level, the TRPV1 cation channel was up-regulated in N30 conditions, while and TRPV1, TRPA1, piezo1 and 2 cation channels, L-type Ca^2+^ channel, and the KCNK2 potassium channel were up-regulated in N90 conditions. The TRP cation channels are typically associated with temperature and pain sensing and can be activated via inflammatory mediators, such as IL-1, IL-6 and ROS;^17, 45, 46^ which we assess in this work. TRP and piezo channels have also been implicated in vibrational mechanotransduction, such as in sterocillia signalling in hearing.^22^ In fact, both TRP and piezo channels have also been linked to cytoskeletal organisation, YES associated protein (YAP, a known osteogenic mechanoregulator^47, 48^) and BMP-2 regulated osteogenesis.^24, 49, 50^ This ties in well with previous findings that 3D osteogenesis can be disrupted with TRPV1 inhibitors^9^ and adds to the body of evidence suggesting that these ion channels have wide ranging effects in cellular mechanotransduction. Again, subtle results observed at N30 were more evident at N90.

The transduction of higher frequency sound waves into biologically relevant signals is a poorly understood mechanism. However, we postulate that cells might convert higher frequency stimuli in constant excitatory responses. For example, at low frequency, Piezo1 behaves like a bandpass filter with a centre frequency at around 10 Hz.^51^ At higher frequencies, the in-phase peak response disappears, and a ‘‘tonic’’ current remains, which in turn increases with frequency. This model predicts that for a 1000 Hz stimulus, a sustained tonic current is expected, which might be comparable with single stimulus peak response when the number of stimulated channels exceeds several hundred.^51^ This behaviour suggests the presence of an underlying molecular-lever mechanism that is able to transduce the mechanical stimulus with a frequency-dependent efficiency. While the specific origin of this mechanism is not yet clear, it is noteworthy that many mechanosensitive ion channels have been found to have common structural features, and it is likely that this is the root for a broad and concerted cell mechanosensitivity.^52^

Untargeted metabolomics analysis led us to look at ROS and inflammation. Subtle increases in ROS and redox-balancing pathways, such as PPP and SOD, were activated in response to nanostimulation at N90 (Figure 3C, Figure 4). Similarly, pro-inflammatory and inflammation-mediating pathways were transcriptionally activated (Figure 5). However, pro-inflammatory cytokines were not detected at the protein level following MSC stimulation at N90 (Figure 5D). Our metabolomic data also reveal hallmarks of increased energy demand during MSC differentiation, which concurs with the literature showing that increased oxidative phosphorylation occurs in MSCs undergoing differentiation to produce more ATP, ^36, 53, 54^. From these findings, we propose that this energy demand drives increased levels of ROS and thus some of the markers of inflammation. It is well known that ROS is generated via increased electron transport chain activity.^55^ However, MSCs counter this increase in potentially damaging pathways.^31^ Inhibiting ROS had only a small effect on osteogenesis (Figure 4E) and likely illustrates that ROS production and inflammation are by-products of osteogenesis rather than drivers of it. Early stage, low level increases in ROS and inflammation are seen in the earliest stages of bone fracture repair – in the inflammatory phase, also known as fracture haematoma formation.^39^ From our data, we speculate that this inflammation is a byproduct of bone cell stimulation. Together, our metabolomic and biochemical data and that already published, lead us to propose a model for nanovibrational stimulation pathways, in which respiration/energy are linked to ROS and inflammation balancing responses and MAPK signalling to drive osteogenesis (Figure 8). However, more work is required to fully elucidate these proposed pathways.

**Figure 8.**
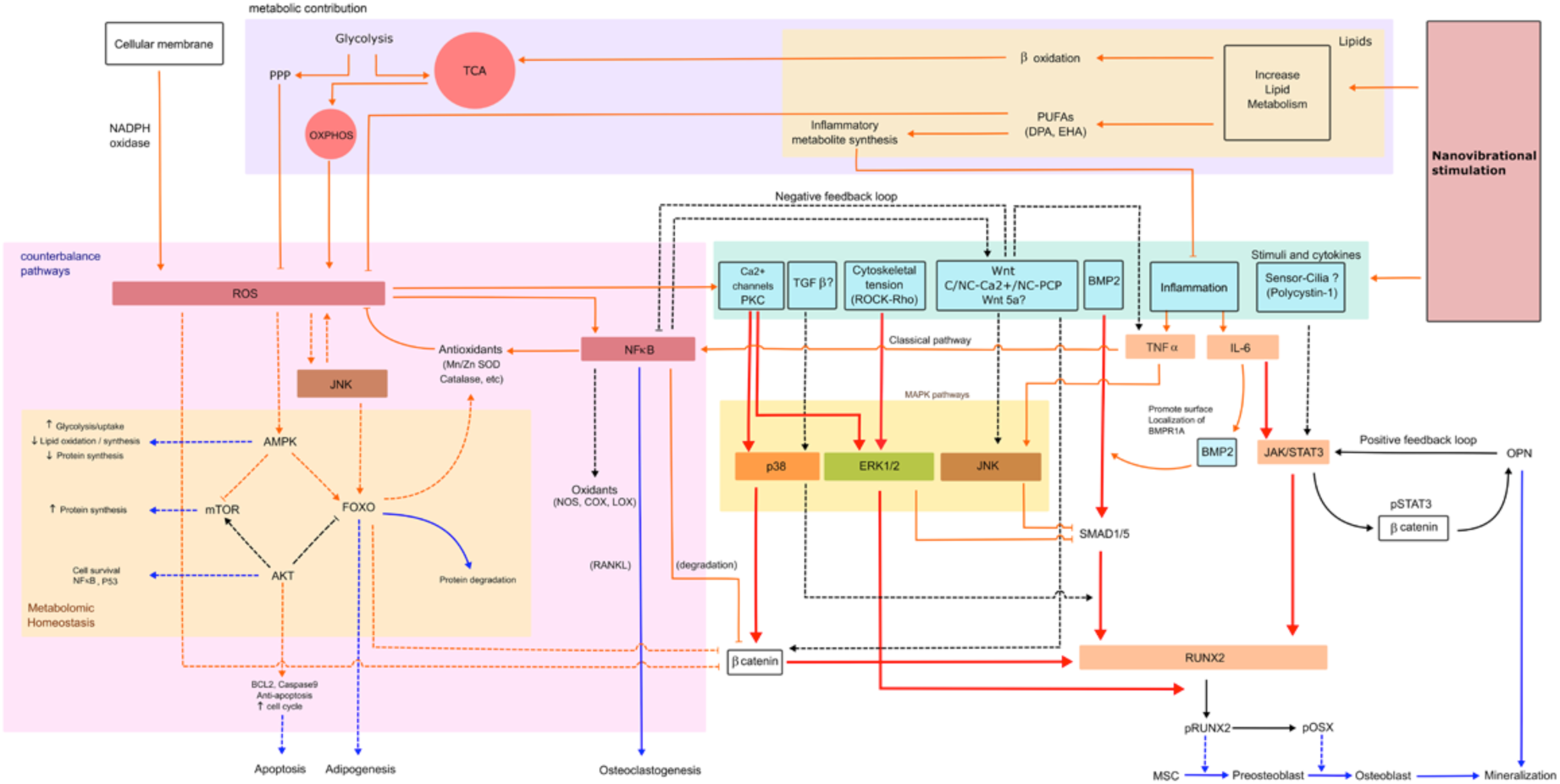
*A model of the mechanisms that contribute to nonvibrational MSC stimulation illustrating that respiration is linked to ROS and inflammation balancing responses and MAPK signalling to drive osteogenesis.* Abbreviations; AMPK (AMP-activated protein kinase), AKT (Protein kinase B (PKB)), BCL2 (B-cell lymphoma 2), C-Wnt (canonical Wnt), COX (cyclooxygenase), FOXO (Forkhead box class O), JAK (Janus kinases), LOX (lipoxygenase), mTOR (mammalian target of rapamycin), NADPH oxidase (nicotinamide adenine dinucleotide phosphate oxidase), NC-Ca^2+^ (noncanonical Wnt/calcium pathway), NC-PCP (noncanonical planar cell polarity pathway), NOS (Nitric oxide synthase), OXPHOS (oxidative phosphorylation), PKC (Protein kinase C), pOSX (phosphorylated osterix), pSTAT3 (signal transducer and activator of transcription protein 3), RANKL (receptor activator of nuclear factor kappa-B ligand), SMAD1/5 (small mothers against decapentaplegic homolog 1/5), Wnt5a (wingless-related integration site). Line description – red line; expected high level expression, orange line; expected low level expression, blue line; phenotype expression, solid line; predicted pathways, dot line; theoretically pathways.

Our data also allow us to speculate that over a threshold amplitude, nanovibrational stimulation could become detrimental to cells as they struggle to balance the increasing levels of ROS and inflammation. Furthermore, our analysis of a non-responding donor MSC line indicates that inflammatory background might be important in the selection of donor material for orthopaedic cell manufacture (supplementary Figure 5).

Finally, our study explored the effects of both an increased N90 amplitude on MSC osteogenesis, and also the generation of a sponge-gel composite that is easier to handle. We assessed MSCs cultured in this gel composite for markers of osteogenesis and inflammation, and observed enhanced osteogenesis in MSCs cultured under N90, relative to N30 conditions, and also some initial expression of IL-6 and NF*κ*B that quickly reduced. This lends further support to the hypothesis that low-level inflammation occurs in nanovibration-enhanced osteogenesis. We also investigated when we could allow the gels to contract onto the sponge to minimise manufacturing time for use in the bioreactor, and found that two weeks of N90 treatment provide the most optimal time.

## Conclusion

In this study, we demonstrate the important contribution of amplitude to the nanovibrational stimulation of osteogenesis in MSCs. We find that an amplitude of N90 produces increased osteogenesis, relative to an amplitude of N30, and use these different responses to investigate subtle changes in adhesion, tension, ion channel regulation, ROS and inflammation in the osteogenic MSC response. Using our bioreactor, we provide new insights into the low-level ROS and inflammatory responses that are typically seen with osteogenesis both in culture, and in the clinic, resultant from the energetic demands of differentiation. We confirm these findings in a newly engineered 3D osteogenic cellular composite, which works with our bioreactor and provides both an enhanced osteogenic environment and handlability.

## Materials and Methods

### Cell culture

Stro-1 selected MSCs from adult human bone marrow (BM) with informed consent from Southampton General Hospital. MSCs were cultured in Dulbecco’s modified essential medium (DMEM; Sigma) supplemented with 10% foetal Bovine Serum (FBS; Sigma), 1% (v/v) L-glutamine (200mM, Gibco), 1% sodium pyruvate (11 mg/ml, Sigma), 1% MEM NEAA (amino acids, Gibco) and 2% antibiotics (6.74 U/mL penicillin-streptomycin, 0.2 µg/mL fungizone; Sigma). MSCs were cultured in an incubator set at 37°C with 5% CO_2_ environment and subcultured to passage 2-3 before use. Culture media was changed every 3 days.

### Hydrogel preparation

0.8 and 1.8 mg/ml collagen hydrogels were prepared using rat tail collagen type I (2 and 5 mg/ml, First link, UK). 10xDMEM (First link, UK) and FBS (Sigma) were added as a cell supplement. 0.1M sodium hydroxide (Fluka, UK) was used for pH titration to achieve pH 7.7-8.0 judged by universal litmus paper and phenol red indicator. After pH titration, Stro-1 selected MSCs were added in to the hydrogel mixture (final cell concentration was 4×10^4^ cells per ml of hydrogel) and decanted as 2.5 ml pre-gels into 6- or 12-well plates. Gelation followed in a 37 °C incubator for 30 min.

### Bioreactor set up

The nanovibrational bioreactor incorporating piezo actuators (Thru-ring actuators; P-010.00H, Physik Instrumente, Germany), was used to stimulate cell cultures. A laptop, as signal generator, was connected to the amplifier (Linn amplifier, Sneaky DS, UK) transferring an electrical signal to the nanovibrational bioreactor causeing piezo expansion. Selected nanovibrational stimulation (NS) frequencies in a .flac file type were operated on Kinsky software (Version 4.3.14) using as a file control panel. To construct the culture plate-bioreactor apparatus, adhesive magnetic sheets (3M, UK) were adhered to the bottom of 6 well- or 12 well-plates (Corning, USA) and attached on the platform of the bioreactor.

### Freeze dried collagen sponge preparation

To prepare the collagen sponges, 5% weight of bovine tendon powder (Collagen Solutions, UK) was used. 0.001 M of HCl at pH 3 was added and then homogenized (TissueRuptor, Qiagen) on ice. The mixed composites were then molded in polystyrene cell culture inserts (0.4 μm Polyethylene terephthalate; PET, membrane, 12 well plate diameter, Greiner bio-one, Austria) and in turn frozen at −80°C for 10 hours. Freeze drying was performed at −110 °C with a vapor pressure at 0 mBar, for 20 hours (VirTis, SP industries, USA). Freeze dried collagen sponges were sterilised by UV light exposure for 1 hour.

### Collagen-hydrogel-sponge composite preparation

To prepare the composites, sponges were placed at the centre of 12 well plates and were weighed down by placing a sterile screw nut on top of the sponges. 1.8 mg/ml collagen hydrogels were prepared and 2.5 ml of hydrogel with MSCs was poured around the sponges. Gelation was allowed at pH 7.7 – 8.0 in a 37 °C incubator. Following the cell-hydrogel gelation, 0.5 ml of secondary hydrogels (1.8 mg/ml) without cells were aliquoted on top the composite to fill any gaps. The composites were then transferred for stimulation on the NS bioreactor. When the stimulation time ended, the composites were detached from the culture wells by spatula allowing contraction before removal for further analysis.

### Scanning electron microscopy (SEM)

The samples were prepared for SEM using critical point drying and sputtering. The samples were mounted onto SEM stubs using double sided conductive tape and silver paint. They were then coated with Gold/ Palladium (approximately 10-20 nm) using a SEM coating system (Q150T ES, Quorum, UK). The samples were viewed on a JEOL6400 SEM running at 10 kV. Porosity diameter was analysed by ImageJ (free download from NIH).

### Composite contraction measurement and time lapse microscopy

In order to monitor the hydrogel contraction for the composites without NS, the percentage of the hydrogel contraction compared to the initial surface area as measured from a top view. The surface areas were measured and analysed by ImageJ software. To evaluate cell migration, composites were cultured for 6 days pre-contraction and 10 days post-contraction. The composites were then imaged using a 10x objective lens at 120 sec intervals for 24 hours at 37°C. Cell velocity and migration directions were analysed using imageJ plugin (manual tracking) and chemotaxis and migration tool (Ibidi).

### Cryosection and immunostaining

After composites were stimulated with NS, the samples were fixed in 4% formadehyde and infused in 30% sucrose in PBS for cryoprotection overnight at 4 °C. The samples were then transferred to embedding moulds with O.C.T. embedding compound (optimal cutting temperature, Tissue-Tek, Sakura) was aliquoted to cover the samples. The samples were then frozen in liquid nitrogen and kept at −80 °C. The frozen samples were sliced into 60 mm thickness sections and attached onto adhesive slides (9597, Tissue-Tek, Sakura, Netherlands). Immunofluorescent staining was then carried out. Samples were rinsed with 1xPBS and fixed with fixative at 37°C for 15 minutes. After that, permeabilisation buffer was added and incubated at 4°C for 5 minutes. Samples were then blocked with 1% BSA in 1xPBS at 37°C for 5 minutes. Primary antibody (P-myosin light chain 2, cell signalling, 3671S, rabbit, 1:50) and rhodamine-phalloidin (Invitrogen, Thermo fisher, 1:500) diluted in 1% BSA in 1xPBS were added and incubated at 37°C for 1 hour. Samples were then washed with 1xPBS/0.5% Tween-20 for 3 times (5 minutes each). Biotinylated secondary antibody (anti-rabbit; Vector Laboratories, USA, 1:50) diluted in 1% BSA in 1xPBS was added and incubated at 37°C for 1 hour. Samples were washed again with 1xPBS/0.5% Tween for 3 times. Streptavidin-FITC (Vector Laboratories, USA, 1:50) diluted in 1% BSA in 1xPBS was incubated at 4°C for 30 minutes. Samples were washed for 3 times. A small drop of DAPI (4’,6-diamidino-2 phenylindole, Vectashield) was placed onto the samples and covered with coverslips. FITC/TRITC channel images were taken (Olympus, US) operated on Surveyor software version 9.0.1.4 (Objective Imaging, UK). Images were processed using ImageJ (Version 1.50g, USA) and Photoshop CS4 (Adobe, version11 extended, Ireland).

### Interferometric measurement

0.8 mg/ml or 1.8 mg/ml collagen hydrogels were prepared. The hydrogel nanovibration was measured by laser interferometric vibrometer (wavelength = 632.8 nm, CW power; 5mW; SIOS, Meβtechnik GmbH, Germany). 3×3 mm of reflective tape was placed on the hydrogels surface underneath 1.5 ml of media to reflect the laser beam. The vibration distance was analysed using INFAS Vibro 1.8.4 software (SIOS, Meβtechnik GmbH, Germany).

### Rheology

To investigate the viscoelastic properties of hydrogels, a modular compact rheometer (MCR 302, Anton Paar, Austria) equipped with parallel plates of 25 mm diameter was used. Measurements were performed at a temperature of 23°C, under a constant normal force and gap size both ranging from 0.1 to 0.3 N and from 1.4 to 2 mm, depending on the samples, respectively. Initial strain sweep tests at a constant frequency of 10 rad/sec were performed to determine the range of linear responce of the hydrogels. Then, the hydrogels’ linear viscoelastic properties were measured by means of frequency sweep tests performed with a constant strain amplitude (of circa 1% for most of the samples) and frequencies ranging from 100 to 0.1 rad/sec.

### AlamarBlue assay

4×10^4^ cells/ml of Stro-1 selected MSCs were prepared in hydrogels or composites and stimulated for 1 and 2 weeks. Samples were washed with warm 1xPBS. 10% (v/v) of AlamarBlue resazurin (Bio-Rad, Watford, UK) diluted in phenol-red free media (D5030, Sigma). After incubation at 37°C and 5% CO_2_ for 5 hours, the supernatant containing the up-taken AlamarBlue was pipetted and transferred into 96 well plates. A microplate reader (Clariostar, BMG Labtech, Germany) was used to detect light absorbance at wavelengths of 570 nm and 600 nm. The percentage of AlamarBlue reduction was calculated as per ^56^.

### Quantitative polymerase chain reaction with reverse transcription (qRT-PCR)

To assess gene expression changes, 4×10^4^ cells/ml of hydrogel were prepared for qRT-PCR. Samples were removed from the bioreactor and transferred to falcon tubes. 2.5 mg/ml of collagenase (Sigma-Aldrich, UK) was added and incubated for 1.5 hours to digest the collagen hydrogels. Trizol (Life Technologies) and chloroform (Sigma-Aldrich, UK) were then added with ratio 5:1 in order to purify nucleic acids. An RNA extraction kit (RNeasy extraction Kit, Qiagen) was used to purify RNA. The concentration of purified RNA was measured by spectrophotometer (Nanodrop 2000c, Thermo scientific, USA). cDNA was then synthesized using the QuantiTect Reverse Transcription Kit (Qiagen). The cycling temperature in each process is shown in Table 1. Forward and reverse primers for qRT-PCR are shown in Table 2. GAPDH, house-keeping gene, was used as internal control of the analysis. SYBR Green dye was used to target synthesized cDNA (Quantifast SYBR Green I, Qiagen). Real time PCR was then performed (7500 Real Time PCR system, Applied Biosystem, USA). 2^−ΔΔCT^ method was used for interpretation^57^.

**Table 1.** Thermal cycler protocol for cDNA synthesis.

| Reaction | Temperature | Times (minutes) |
| --- | --- | --- |
| Genomic DNA elimination | 42°C | 2 |
| Reverse transcription | 42°C | 30 |
| Inactivation | 95°C | 3 |

**Table 2.**
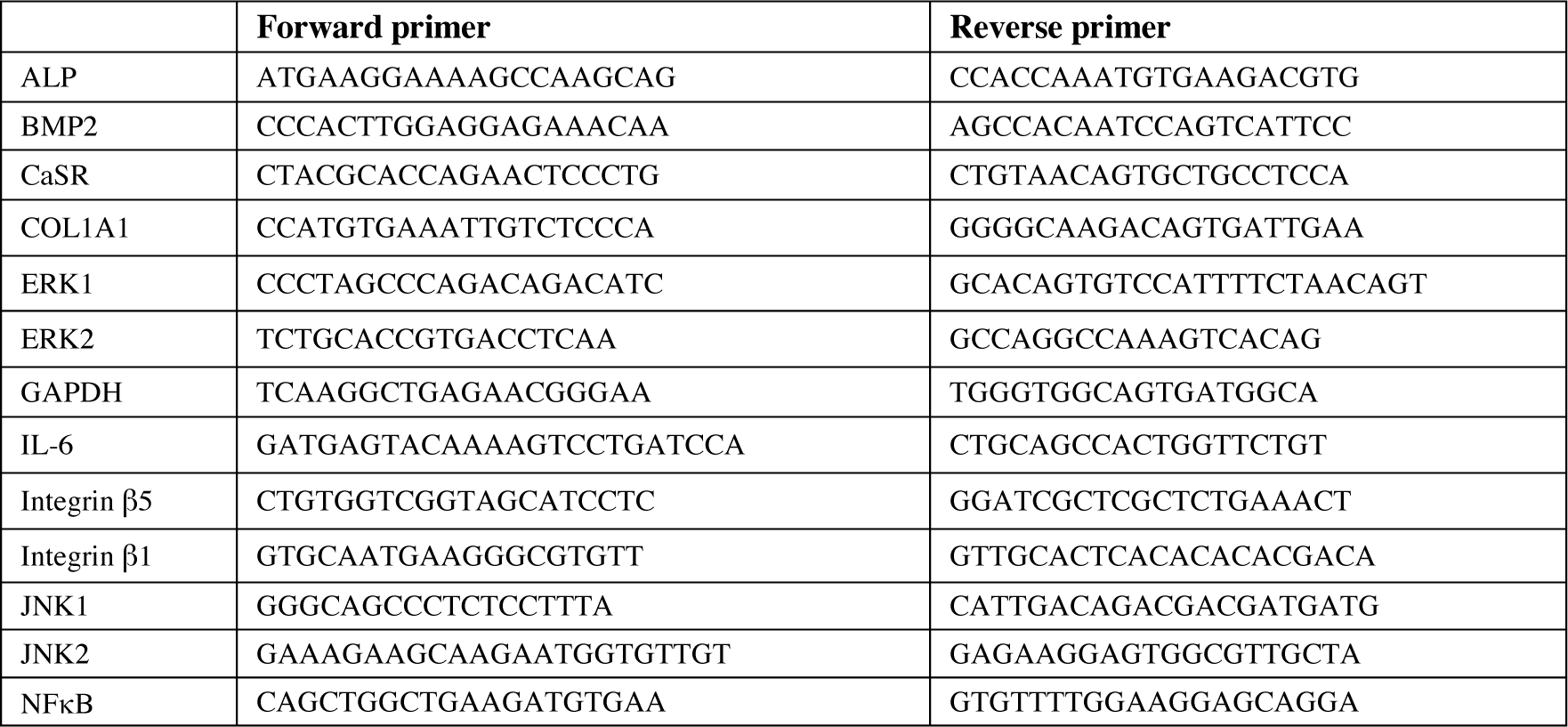
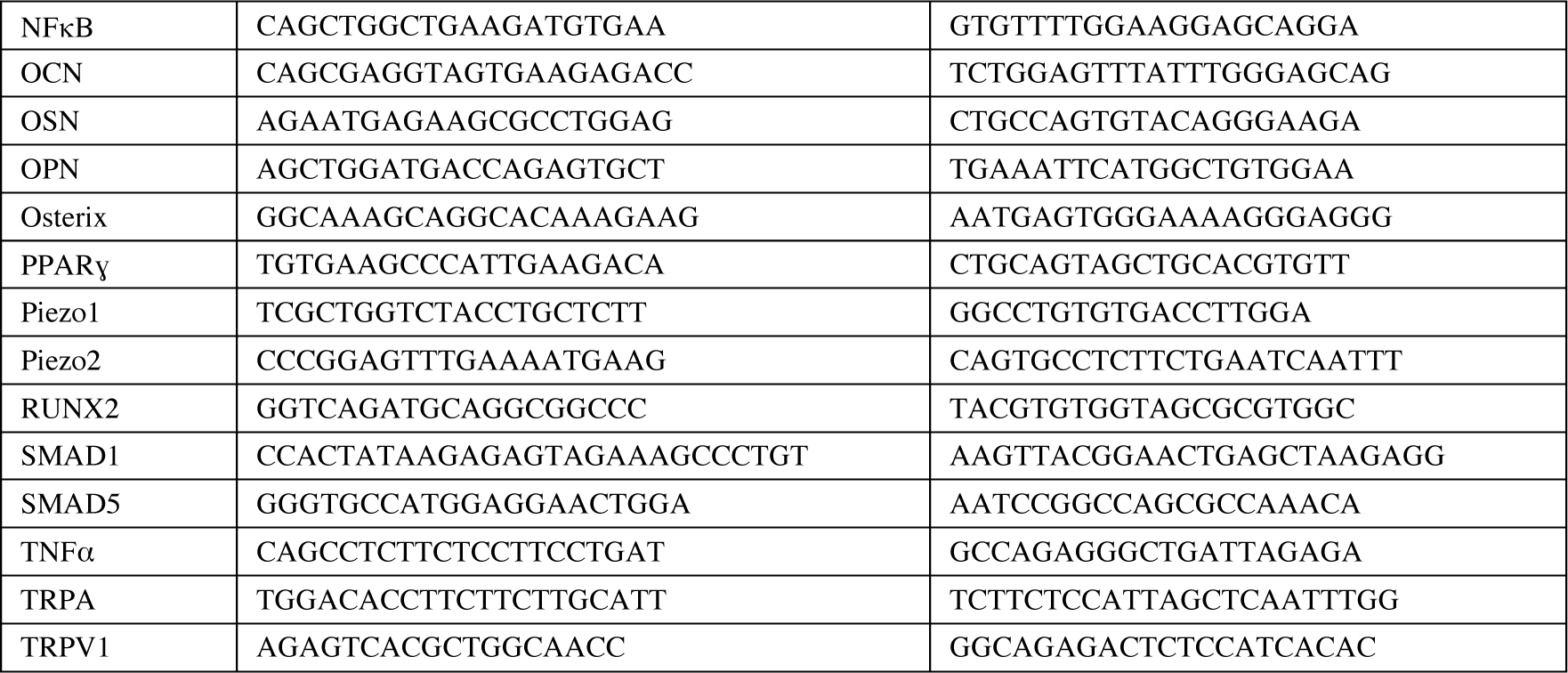
Primer sequences used in qRT-PCR.

### Inhibitor studies

4×10^4^ cells/ml of Stro1 selected MSCs in 1.8 mg/ml collagen hydrogels were stimulated with N30 and N90 for 9 days. Inhibitors, which were diluted in basal media to working concentration, were added at day 2 (list of inhibitors and used concentration are shown in Table 3). Culture media with diluted inhibitor was changed every 2 days.

**Table 3.** List of inhibitors and working concentration.

| Function | Inhibitors | Cat No./Batch | Working concentration |
| --- | --- | --- | --- |
| ERK inhibitor | U126, Tocris | 1144/5 | 10 μM |
| P38 inhibitor | SB 202190, Tocris | 1264/5 | 5 μM |
| JNK inhibitor | SP600125, Tocris | 1496/10 | 25 μM |
| NFκB inhibitor | TPCA-1, Tocris | 2559/5 | 5 μM |
| TNF alpha | R 7050, Tocris | 5432/1 | 2 μM |
| ROCK inhibitor | Y-27632, Tocris | 1254/35 | 10 μM |
| ROS inhibitor | N-acetyl cysteine, Sigma Aldrich | WXBC4028V | 10 μM |

### Protein antibody microarrays

To manufacture the arrays, commercial antibodies (Table 4) were buffer exchanged into PBS and quantified by bicinchoninic acid (BCA) assay. Antibodies were diluted to print concentration in PBS and printed in six replicates on Nexterion H amine reactive hydrogel coated glass slides (Schott AG, Mainz, Germany) using a SciFLEXARRAYER S3 piezoelectric printer (Scienion, Berlin, Germany) under constant humidity (62% +/− 2%) at 20 °C. Each feature was printed using ≈1 nL of diluted antibody via an uncoated 90 µm glass nozzle with eight replicated subarrays per microarray slide. After printing, slides were incubated in a humidity chamber overnight at room temperature to facilitate complete conjugation. The slides were then blocked in 100 × 10^−3^ M ethanolamine in 50 × 10^−3^ M sodium borate, pH 8.0, for 1 h at room temperature. Slides were washed in PBS with 0.05% Tween 20 (PBS-T) three times for 2 min each wash followed by one wash in PBS, dried by centrifugation (470 × g, 5 min), and then stored with desiccant at 4 °C until use; antibody microarrays were verified to remain active for at least 2 weeks after printing and all incubations were carried out within that timeframe.

**Table 4.**
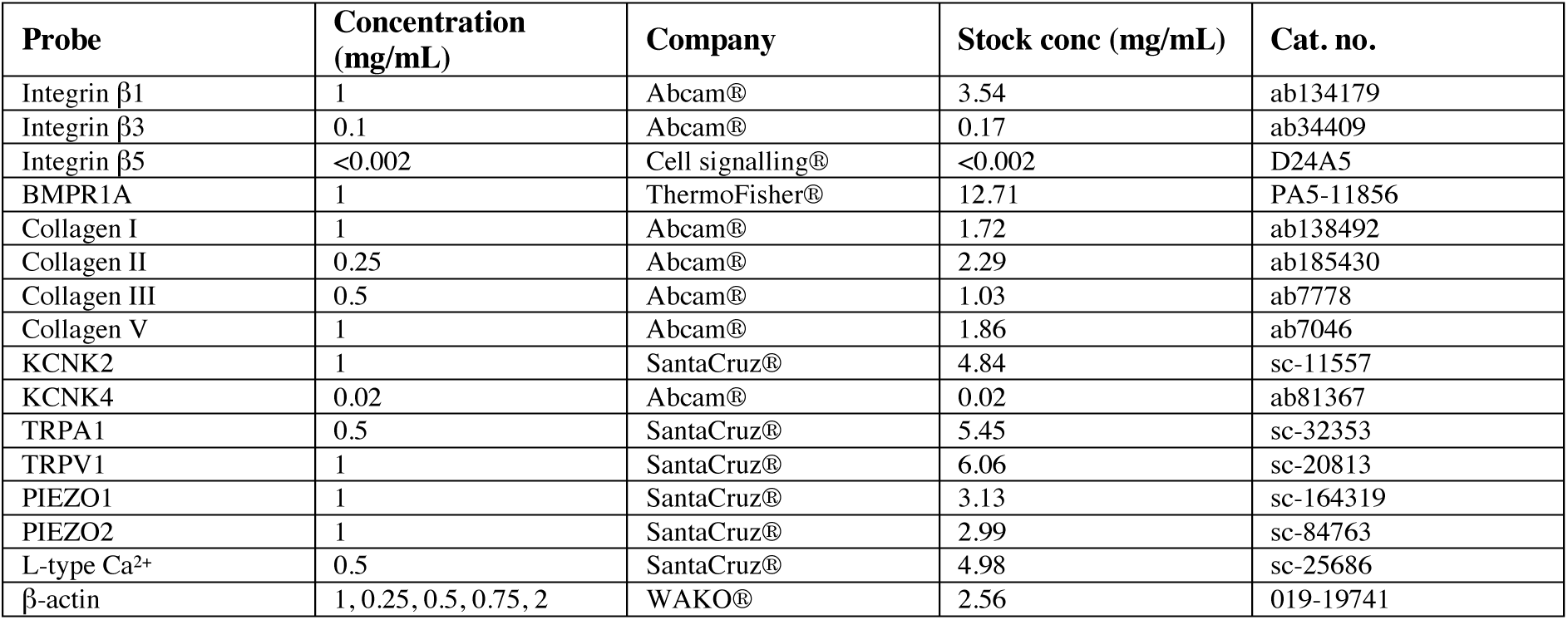
List of commercial antibodies used in the protein antibody microarray.

4×10^4^ cells/ml of Stro-1 selected MSCs in 1.8 mg/ml collagen hydrogels were stimulated with N30 and N90. After stimulation for 1 and 2 weeks, hydrogel samples were digested with collagenase. Total protein was quantified using micro-BCA kit (Pierce, Thermo Fisher). Initially, one labeled sample was titrated (2.5–15 µg mL^−1^) for optimal signal to noise ratio and all samples were subsequently incubated for 1 h at 23 °C at 9 µg mL−1 in Tris-buffered saline (TBS; 20 × 10^−3^ m Tris-HCl, 100 × 10^−3^ m NaCl, 1 × 10−3 m CaCl_2_, 1 × 10^−3^ m MgCl2, pH 7.2) with 0.05% Tween 20 (TBS-T). All microarray experiments were carried out using three replicate slides. Alexa Fluor 555 labeled MSC lysates (10 µg mL^−1^) were incubated in two separate subarrays on every slide to confirm retained antibody performance and printing, respectively. After incubation, slides were washed three times in TBS-T for 2 min per wash, once in TBS and then centrifuged dry. Dried slides were scanned immediately on an Agilent G2505 microarray scanner using the Cy3 channel (532 nm excitation, 90% photomultiplier tubes (PMT), 5 µm resolution) and intensity data were saved as .tif files. Data were normalized to the mean of three replicate microarray slides (subarray-by-subarray using subarray total intensity, n = 3, 18 data points). β-actin was used as internal protein control. Heatmaps were generated by Hierarchical Clustering Explorer v3.0.

### Metabolomics

Stro-1-selected MSCs seeded with 4 x10^4^ cells/ml density in 1.8 mg/ml collagen hydrogels were stimulated with N30 and N90. After 1 and 2 weeks NS the gels were homogenized on ice and metabolites were then extracted using a chloroform/methanol/water (1:3:1 ratio) extraction buffer. Samples were agitated on a shaker at 4°C for 1 hour and in turn centrifuged at 13,000g at 4°C for 5 minutes. Hydrophilic interaction liquid chromatography-mass spectrometry was performed (Dionex, UltiMate 3000 RSCL system, Thermo Fisher Scientific, Hemel Hempstead, UK) using a ZIC-pHILIC column (150 mm × 4.6 mm, 5 μm. column, Merck Sequant). The datasets were processed using XCMS (peak picking), MzMatch (filter and grouping), IDEOM (post processing filtering and identification). Metaboanalyst was used to generate heatmaps and PCA analysis. KEGG database and Ingenuity Pathway Analysis (IPA) software were used for metabolomic pathway analysis.

### Reactive oxygen species measurement

Stro-1-selected MSCs seeded with 4 x10^4^ cells/ml density in 1.8 mg/ml collagen hydrogels were stimulated with N30 and N90 for 7 days. The samples were incubated in 2.5 mg/ml collagenase for an hour and were then centrifuged at 200xg for 4 minutes. Following that, the cell pellets were incubated for an hour in 2 uM 2’,7’-dichlorodihydrogen-fluorescein diacetate (H_2_DCF-DA, Invitrogen) in phenol red free media (Sigma, D5030). In the positive control group, 500 uM hydrogen peroxide was added. After incubation, the samples were then centrifuged and resuspended in 250 ul of flow cytometry buffer (2% FBS, 2 mM EDTA in 1xPBS) and transferred to 96 well plates. Resuspended MSCs in 96 well plates were incubated for 30 minutes. Signal of H_2_DCF-DA fluorescein was detected by using flow cytometry at 492-295 nm for excitation and 517-527 nm for emission.

### ELISA of interleukin-1β

Stro-1-selected MSCs seeded with 4 x10^4^ cells/ml density in 1.8 mg/ml collagen hydrogels were nanostimulated for 7 days. Hydrogels were digested with collagenase (Sigma-Aldrich, UK). Protein was extracted using RIPA lysis buffer containing phosphatase and protease inhibitors. Total protein concentration was quantified using BCA kits (Pierce, ThermoFisher). Human IL-1β kits (DY201-05, R&D systems) were used for analysis. Working concentration of reagents (Human IL-1*β* capture antibody; 840168, Human IL-1*β* detection Antibody; 840169, Human IL-1*β* standard; 840170, Streptavidin-HRP; 893975) were prepared as per manufacturer’s instructions. To coat the captured antibody onto the ELISA plate, 100 μl captured antibody was added into 96-well plate and incubated overnight at 4°C. The plate was then washed with washing buffer (0.05% Tween 20 in 1xPBS (pH 7.4) 3 times using multichannel pipettes. 300 μl of reagent diluent (Reagent diluent concentrate 2, DY995, R&D systems) was added and incubated for an hour to block the coated plate. 100 μl of samples and of standards in reagent diluent (Human IL-1*β*, 840170) were added and incubated for 2 hours at room temperature. Samples were washed 3 times with washing buffer. 100 μl of streptavidin-HRP was added and incubated for 30 minutes at room temperature. Samples were washed 3 times. 100 μl substrate solution (1:1 of colour reagent A; H_2_O_2_ and colour reagent B; tetramethylbenzidine) wereadded and incubated for 20 minutes. 50 μl stop solution (2NH_2_SO_4_, DY994, R&D systems) was added. A microplate reader was used to determine the optical density at 450 nm and 570 nm.

### Statistics

To compare the means of samples of more than two groups, one-way ANOVA with a Tukey post hoc test was used in qRT-PCR and AlamarBlue assays. Two-way ANOVA with Tukey post hoc test was used to analyse hydrogel and composites contraction by times. To compare the data between two groups, two tailed, paired t-test were used for AlamarBlue Assay of composites. Two tailed, Mann-Whitney U tests were used in interferometric measurement and qRT-PCR. Biological sample populations with 4 replicates were always used. All results are shown in mean ± standard deviation with 95%, 99% and 99.9% of accuracy (* P≤ 0.5, ** P≤ 0.01, *** P≤ 0.001). Replicate detail for each experiment is shown in Table 5.

**Table 5.**
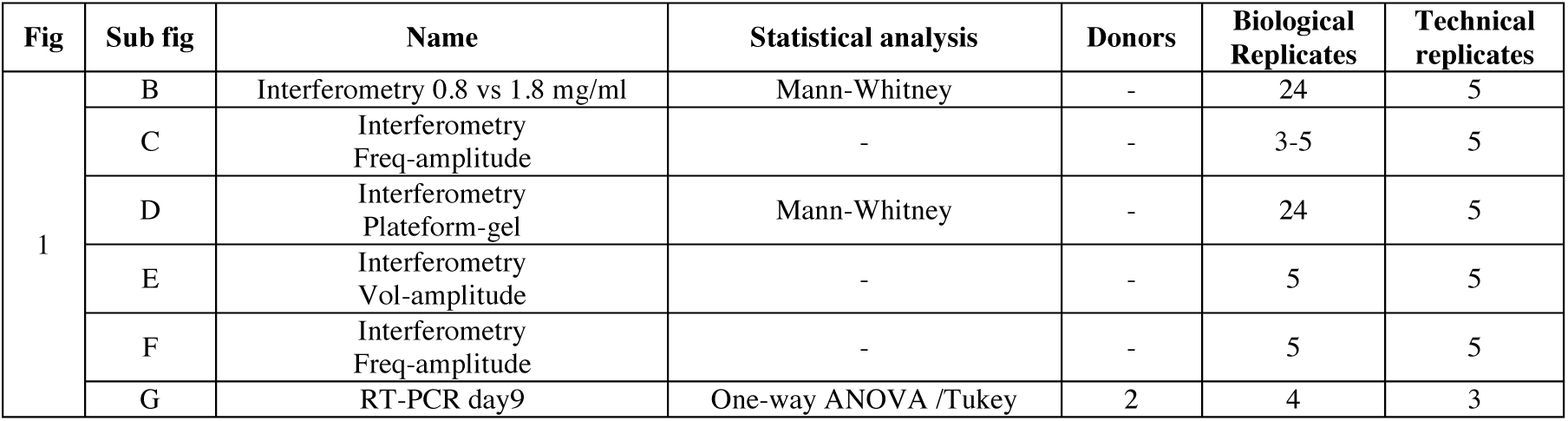
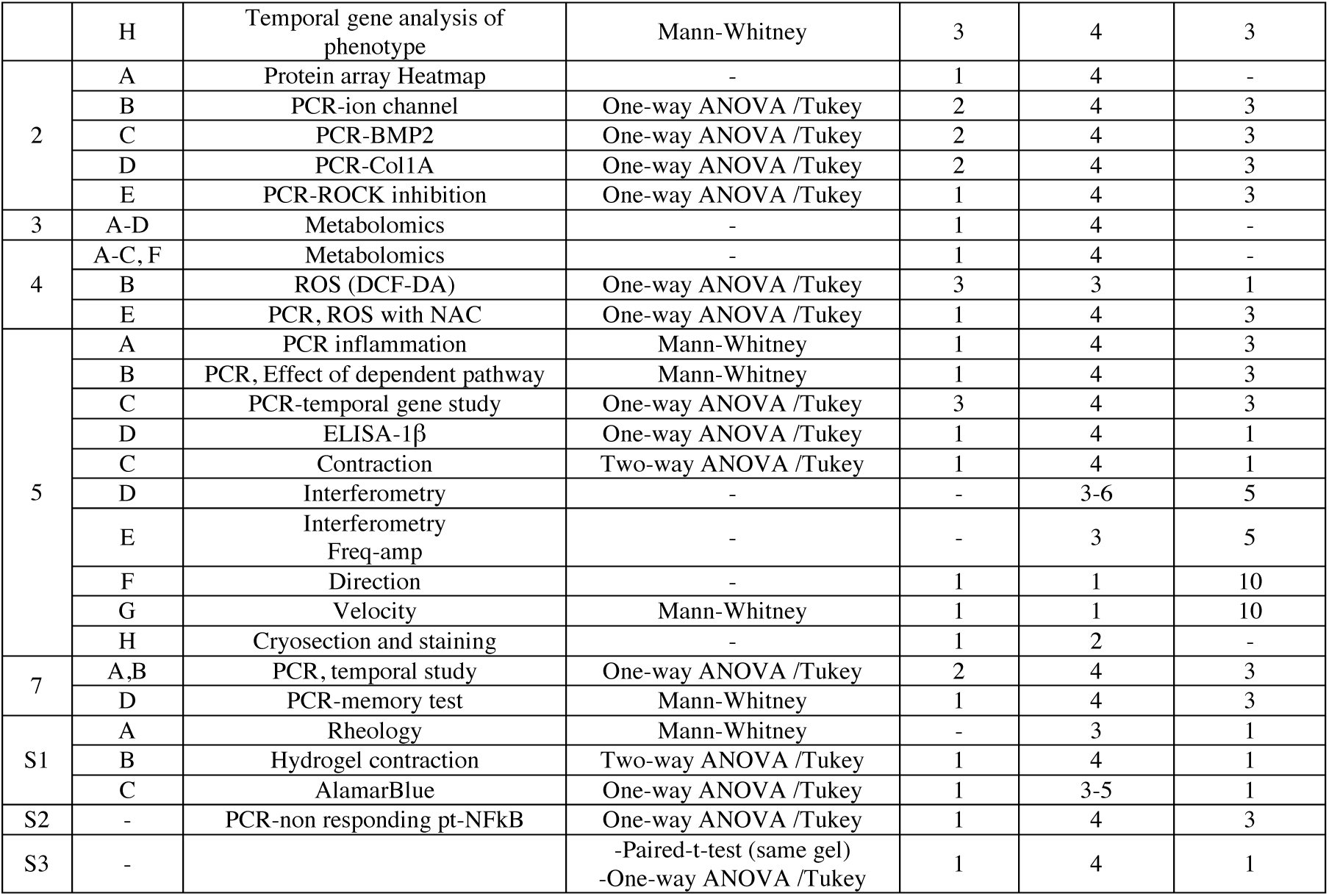
Details of sample replicates and statistical tests used in figures.

## Author Contributions

WO, PMT, MJD, KET, MJPB, SR, MS-S, ROCO conceived the experiments. WO, PMT, KB, MF-Y, KET, PGC, PC performed the experiments. JW, ROCO, MF-YWO, RMDM, PC, SR, MS-S provided materials and expertise. PMT, MJD, PC analysed the data. MV, RMDM, ROCO provided critique and context for the data. WO, PMT, MJD wrote the manuscript. WO, MJD prepared the figures. All authors read and commented on the manuscript.

## Acknowledgement

W.O. was supported by a scholarship from the Royal Thai Government and Faculty of medicine, Prince of Songkla University. The work was supported by grants to M.J.D. from BBSRC (BB/P00220X/1, BB/S018808/1) and EPSRC (EP/N013905/1, EP/P001114/1) and to M.J.P.B from Science Foundation Ireland (16/BBSRC/3317). We thank Mrs Carol-Anne Smith for technical assistance. We thank Jane Alfred, PhD, from Catalyst Editorial (www.catalyst-editorial.co.uk) for editing a draft of this manuscript.

## Supporting Information

**Supplementary Figure 1.**
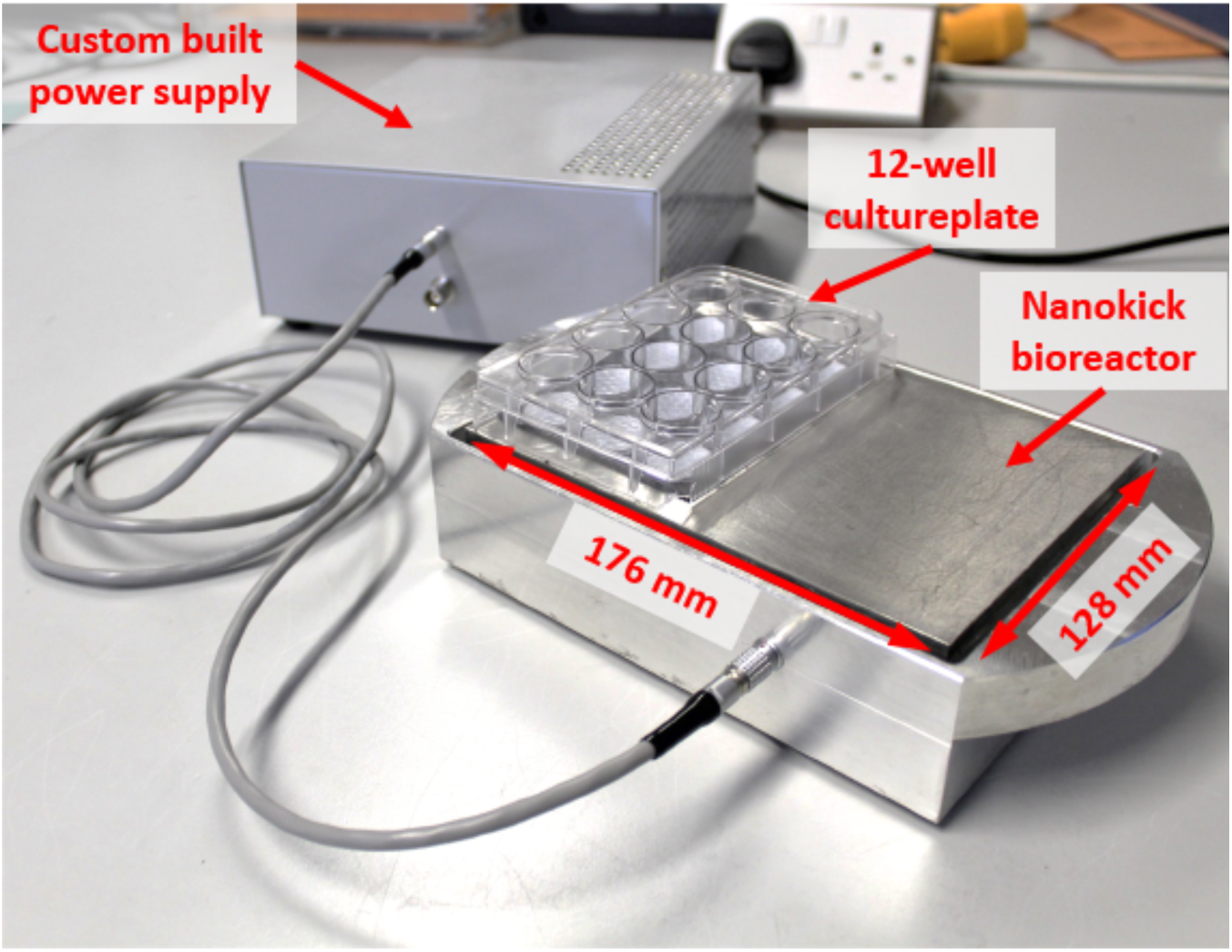
Nanovibrational bioreactor. The Nanovibrational bioreactor system comprises a custom power supply unit, which delivers a 1 kHz sine wave signal to a piezo array under the top plate, causing them to oscillate with an amplitude of 30 nm. The stainless steel top plate can accommodate 2 standard multiwell culture plates or a single T-150 flask attached via rubber magnets.

**Supplementary figure 2.**
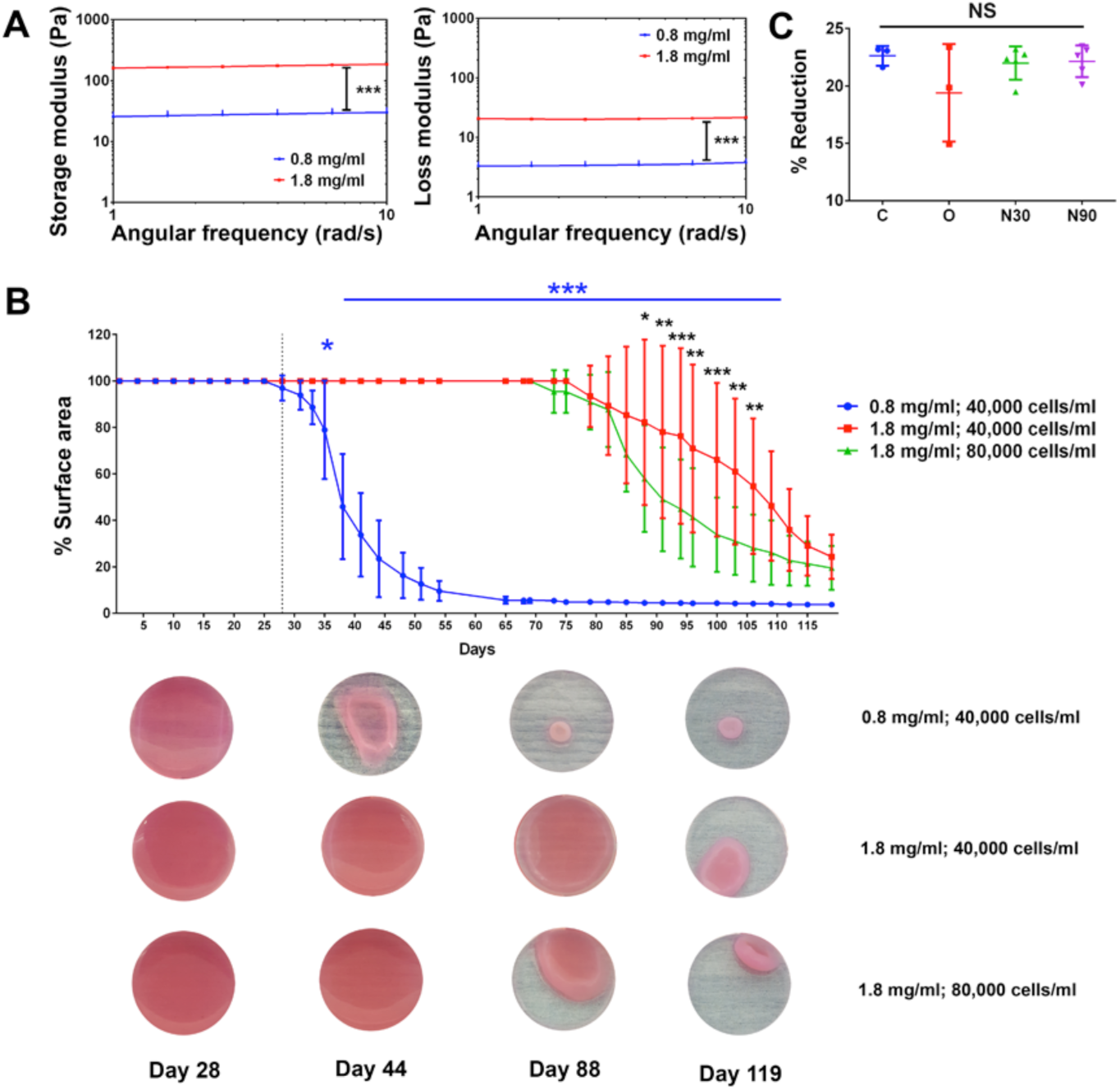
Rheology and contraction characterization. (A) Rheology data for 0.8 and 1.8 mg/ml collagen gels (n=3). (B) Temporal culture data showing that use of the higher-concentration gels prevents possible early contraction (n=4, note that images are of 13 mm diameter, 24 well plates)). (C) Alamar blue assay showing lack of cytotoxic effects for control, OGM (osteogenic media) and nanovibrational stimulation (d=1, r=3-5, t=1, stats by ANOVA with Tukey test).

**Supplementary figure 3.**
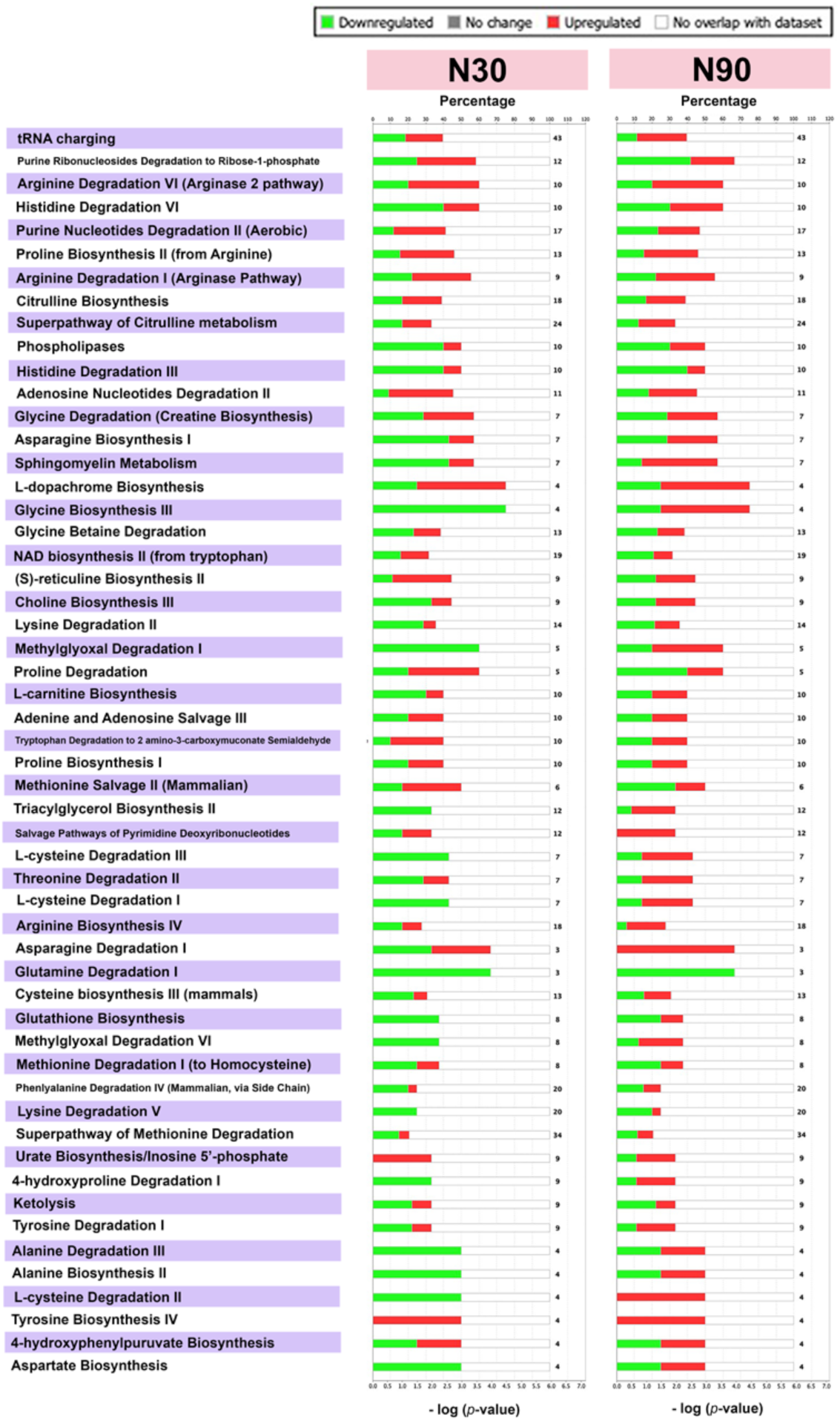
Day 7 metabolomics data for N30 and N90 stimulation. Ingenuity pathway analysis of untargeted metabolomics data derived from MSCs after 7 days of culture with N30 or N90 nanostimulation vs control conditions reveals a preponderance of up-regulated pathways (shown in red) for the N30 condition, which were further upregulated in the N90 condition (down-regulations depicted in green, d=1, r=4, t=1).

**Supplementary figure 4.**
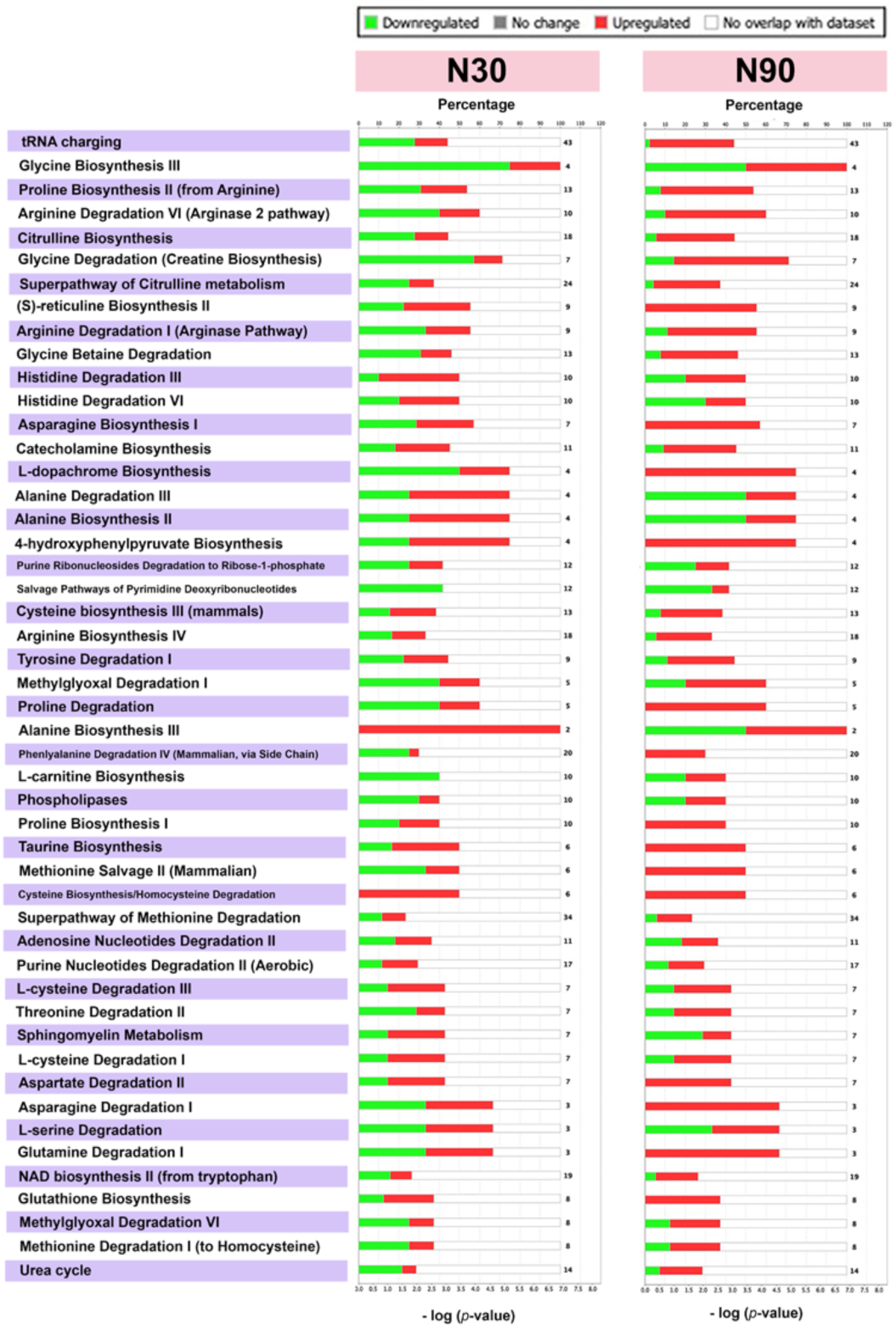
Day 14 metabolomics data for N30 and N90 stimulation. Ingenuity pathway analysis of untargeted metabolomics data derived from MSCs after 14 days of culture with N30 or N90 nanostimulation vs control conditions, shows most pathways are up-regulated (shown in red) for both conditions, but most notably for N90. An increase in pathway down-regulation (shown in green) was also noted compared to day 7 data (d=1, r=4, t=1).

**Supplementary figure 5.**
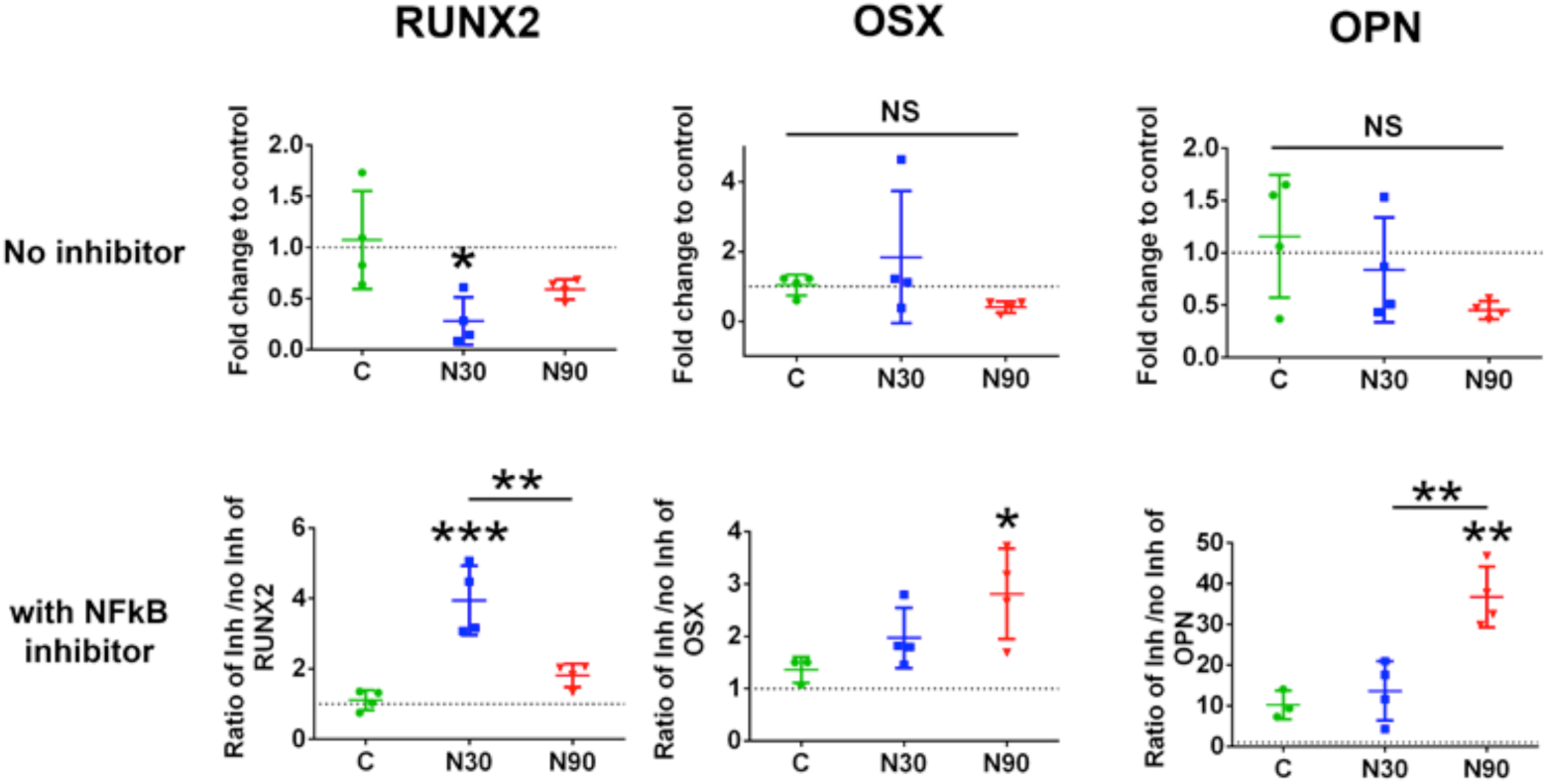
Testing non-responing donor cells. Osteogenic marker expression (RUNX2, OSX, OPN) in MSCs cultured with/without NFκB inhibitor in control, N30 and N90 culture conditions for 9 days with non-responding MSCs (d=1, r=4, t=3), as assessed by qPCR. Error bars represent mean ± SD, significance calculated using ANOVA and Tukey test where *=p<0.05, **=p<0.01, ***=p<0.001).

